# Excitable Rho dynamics drive cell contractions by sequentially inducing ERM protein-mediated actin-membrane attachment and actomyosin contractility

**DOI:** 10.1101/2023.12.19.572346

**Authors:** Seph Marshall-Burghardt, Rodrigo A. Migueles-Ramírez, Qiyao Lin, Nada El Baba, Rayan Saada, Mustakim Umar, Arnold Hayer

## Abstract

Migration of endothelial and many other cells requires spatiotemporal regulation of protrusive and contractile cytoskeletal rearrangements that drive local cell shape changes. Unexpectedly, the small GTPase Rho, a crucial regulator of cell movement, has been reported to be active in both local cell protrusions and retractions, raising the question of how Rho activity can coordinate cell migration. Here we show that Rho activity is absent in local protrusions and active during retractions. During retractions, Rho rapidly activated ezrin-radixin-moesin proteins (ERMs) to increase actin-membrane attachment, and, with a delay, non-muscle myosin II (NMII). Rho activity was excitable, with NMII acting as a slow negative feedback regulator. Strikingly, inhibition of SLK/LOK kinases, through which Rho activates ERMs, caused elongated cell morphologies, impaired Rho-induced cell contractions, and reverted Rho-induced blebbing. Together, our study demonstrates that Rho activity drives retractions by sequentially enhancing ERM-mediated actin-membrane attachment for force transmission and NMII-dependent contractility.

## Introduction

In response to external stimuli, cells can polarize for directed migration, forming distinct cytoskeletal structures specific to the cell front and back. In the absence of directional cues, many adherent cell types lose their front-back polarization, but remain motile and undergo cycles of random protrusion and retraction, often occurring in a wave-like manner^1–4^. This intrinsic cellular behavior occurs on the timescale of minutes and is also widely observed in migrating cells, both in 2D and in 3D environments^5–9^. The small GTPases of the Rho family (RhoGTPases), in particular the Rac, Cdc42, and Rho subfamilies (referred to as Rac, Cdc42, and Rho hereafter), regulate actin cytoskeletal dynamics in motile cells by driving protrusive and contractile cell shape changes. Across many cell types and contexts, Rac and Cdc42 have well-defined roles in cellular protrusions, where they promote actin polymerization-driven cell edge extensions^10–15^. Although widely accepted as the master regulator of cellular contractility through its effectors Rho associated kinase (ROCK) and non-muscle myosin II (NMII)^16,17^, Rho ‘s role in regulating cell motility remains incompletely understood. Elevated Rho activity has been observed in cell retractions^18–22^, but also near the cell edge in protrusions and ruffles, where it showed high correlations with local outward cell edge movements^14,23–27^. A plausible effector downstream of Rho that drives protrusions is the formin mDia1^14,28^. However exogenous activation of Rho generally causes cells to contract and subcellular activation of Rho using optogenetics has not yet been observed to induce protrusions^29–33^. In addition to the established Rho-activated targets ROCK and mDia1, Rho can activate the ezrin-radixin-moesin family proteins (ERMs), through the kinases SLK (Ste20-like kinase) and LOK (lymphocyte-oriented kinase), which have recently emerged as Rho effectors^34–37^. SLK/LOK activate ERMs^35,38^, which, when activated, link actin to the plasma membrane, to form the actin cortex and therefore control shape and mechanical properties of cells^39^. The actin cortex is involved in cell motility in multiple ways. A contractile cortex can drive cell rear retraction or generate pressure during amoeboid cell migration^40,41^. In protrusions, the actin cortex forms a barrier that has to locally weaken or detach from the plasma membrane before protrusions can be initiated^42,43^.

We set out to determine whether Rho activity drives protrusions or retractions and to test for a possible involvement of ERM activation downstream of Rho. We visualized Rho activity along with the Rho effectors NMII or ERMs using fluorescence-based reporters in single unpolarized endothelial cells that exhibit random cycles of minute and µm-scale protrusions and retractions. We found Rho was enriched in retractions, but not in protrusions, and identified that edge-proximal Rho dynamics were pulsatile, displaying hallmark characteristics of excitability. Rho activation led to the sequential (i) SLK/LOK mediated activation of ERMs to enhance actin-membrane attachment and (ii) ROCK-dependent NMII activation to generate contraction. Activated ERMs coincided with Rho in retractions, whereas NMII accumulated with a delay. The inhibition of SLK/LOK kinases resulted in elongated cell phenotypes and impaired Rho-induced cell contractility, demonstrating that both ERMs and NMII are required for contractile cell shape changes downstream of Rho.

## Results

### Sparsely plated HT-HUVEC exhibit, random, minute-scale protrusion-retraction cycles

In the absence of directional cues, HT-HUVEC (hTERT immortalized human umbilical vein endothelial cells^31^) exhibit random motile behavior when plated on uniform substrates at low density. Such random motility is characterized by cycles of protrusions and retractions that occur on a timescale of minutes (Fig. 1a, Video 1). We found that such behavior was particularly suitable for quantitative analysis when observed within a few hours after cell plating, when cells had completed spreading and assumed relatively homogenous and compact adhesion surfaces while showing active protrusion/retraction behavior. We quantified cell shape changes based on time-lapse sequences of cells stably expressing fluorescence-based reporters, automated cell segmentation, followed by edge velocity tracking, using established methods^14,19^. For cell edge velocity tracking, segmented cell outlines were divided into 180 equally spaced peripheral coordinate windows spanning the entire cell perimeter. Each window had an adjustable depth towards the cell interior in which fluorescencebased reporter activity could be measured (Fig. 1b). Window velocities were calculated using per frame displacements, resulting in 2D spatiotemporal edge velocity maps (Fig. 1b,c). From such maps, we identified protrusions and retractions as events exceeding defined minima of outward and inward velocity and spatiotemporal scale (Fig. 1c, Materials and Methods). This analysis revealed that over a 1h duration, randomly sampled cells (n=55) had on average 27.63 ± 10.27 (mean±SD) individual protrusion events, each with an average mean velocity of 6.39 ± 0.61µm/min, lasting for 4.21 (± 0.75) min, and spanning 8.26% (± 1.32%) of the cell perimeter. In comparison, the same cells had 20.79 ± 12.17 retraction events with a mean velocity of -5.93 ± 0.63µm/min, lasting for 5.03 (± 1.10) min and spanning 6.69% (± 1.10%) of the cell perimeter (Fig. 1d).

**Figure 1.**
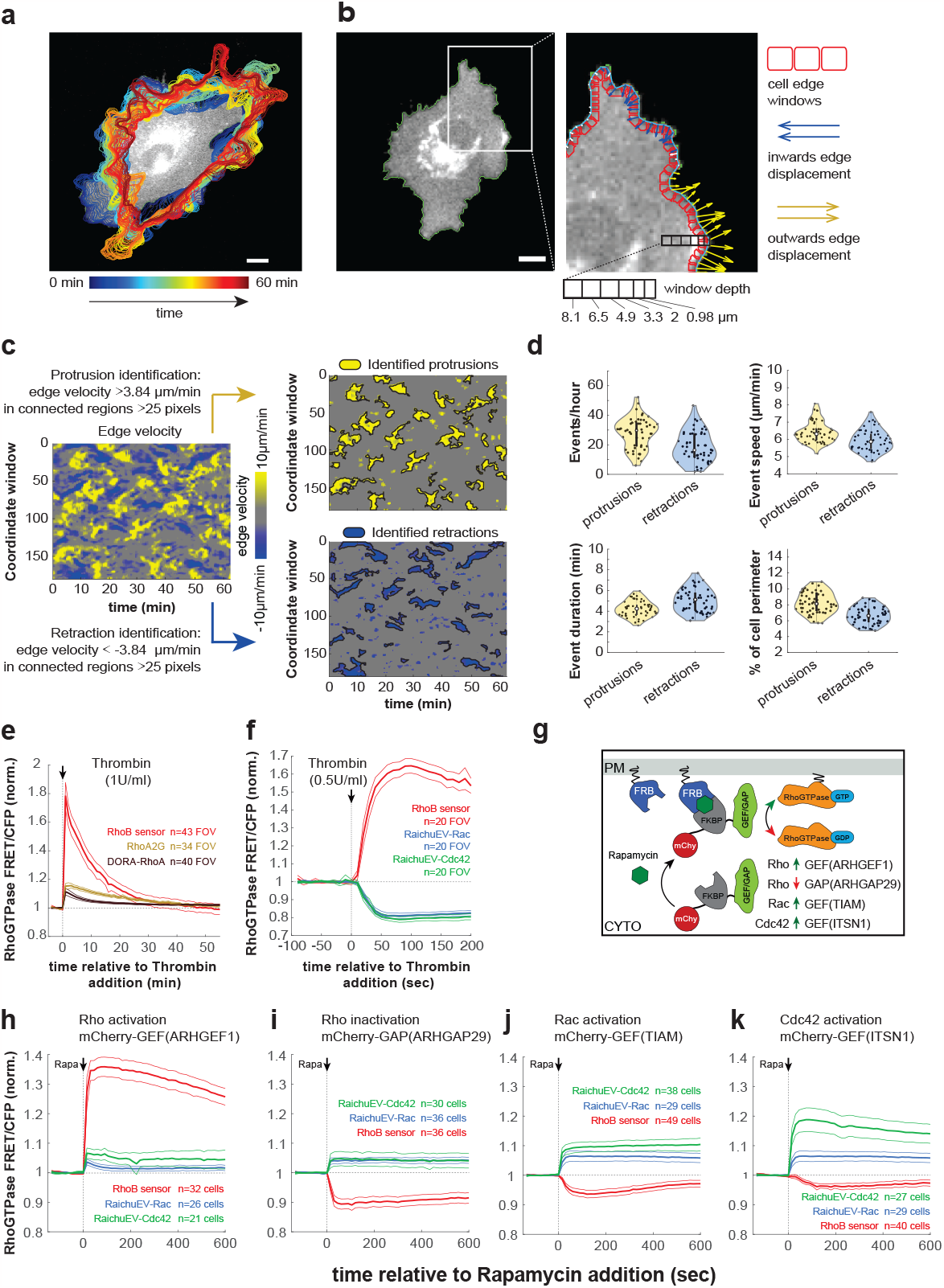
Random motility of HT-HUVEC, cell edge velocity tracking, validation of FRET probes for Rho, Rac, and Cdc42 in HT-HUVEC. (**a)** HT-HUVEC cell expressing the RhoB sensor, overlayed with temporally color-coded cell outlines to illustrate protrusion retraction cycles over 60min. Scale bar, 10µm. (Video 1). **(b)** Illustration of cell edge velocity analysis (Materials & Methods). *(Left)* Masked HT-HUVEC cell expressing the RhoB sensor. Scale bar 10 µm. *(Right)* close-up showing equally spaced cell edge windows (red) used for analysis and their movements (blue, yellow arrows). **(c)** *(Left)* Corresponding cell edge velocity map from (a), *(top right)* automatically segmented protrusions, and *(bottom right)* retractions, based on defined spatiotemporal parameters (Materials & Methods). **(d)** Spatiotemporal parameters of individual protrusions and retractions as identified in (c). Data points are individual events, violin plot displays means ± 25th/75th percentile of events as bolded black lines from n=54 cells, from 3 biological replicates. **(e)** FRET probes for Rho (RhoB sensor, DORA-RhoA, RhoA2G) were stably expressed in HT-HUVEC and their responses to 1U/ml thrombin were measured over 55min in sub-confluent cultures. The RhoB sensor showed by far the strongest response. Responses were normalized to control-treated cells, means of n= number of fields of view analyzed as indicated ± 95% CI, from 3 biological replicates. **(f)** The responses of the RhoB sensor and FRET probes for Rac and Cdc42 (RaichuEV-Rac and RaichuEV-Cdc42) to thrombin stimulation (0.5U/ml) were assessed, similar to (e). Means of n=20 fields of view per condition ± 95% CI, from 2 biological replicates. **(g)** Strategy to acutely (in)activate RhoGT-Pases by plasma membrane translocation of RhoGEF and RhoGAP domains through addition of rapamycin. **(h-k)** HT-HUVEC expressing RhoB sensor, RaichuEV-Rac, or RaichuEV-Cdc42 were transiently contransfected with Lyn-FRB and mCherry-FKBP-RhoGEF/GAP as indicated, and RhoGTPase FRET/CFP signals were measured in response to rapamycin addition (0.5µM). Means ± 95% CI, normalized to untransfected cells, from n=number cells as indicated, from 2 biological replicates.

### Spatiotemporal analysis of Rho, Rac, and Cdc42 using FRET-based activity probes in randomly motile HT-HUVEC

To enable spatiotemporal analysis of Rac, Cdc42, and Rho activities in thresholded protrusions and retractions, we first tested the suitability of existing FRET-based activity probes in HT-HUVEC. For Rho, we tested RhoA-based RhoA2G^44^, DORA-RhoA^25^, and a RhoB-based RhoB sensor^20^. The probes were stably expressed in HT-HUVEC via lentiviral transduction and their responses to acute, pathway-selective perturbations were assessed (Materials and Methods). Activation of Rho using thrombin resulted in increased Rho activity as reported by all three probes, with the RhoB sensor showing by far the strongest response (Fig. 1e,f). RhoA and RhoB share >85% sequence identity, and functionally many GEFs, GAPs, and downstream effectors^34,45^. RhoA and RhoB differ in their C-terminal domains, resulting in distinct lipid modifications and localization patterns, with RhoA and RhoB being mainly cytoplasmic and membrane-localized, respectively^46,47^. Consistent with this, the RhoA-based probes RhoA2G and DORA-RhoA primarily localized to the cytoplasm, whereas the RhoB sensor was mainly plasma membrane-localized (Supplementary Fig. 1a). We found that local Rho activities reported by RhoA2G and DORA-RhoA appeared to be affected by the local cell geometry, when viewed by epifluorescence microscopy. The highest activities were generally enriched in the cell periphery and lowest activities in the cell center, corresponding to high and low membrane/cytoplasm ratios, respectively (Supplementary Fig. 1a (i,ii)). To assess a possible localization bias of reported Rho activities, we performed pixel-by-pixel correlations between normalized Rho activities and their localizations for each of the three probes. RhoA2G and DORA-RhoA both yielded significantly stronger anticorrelations compared to the RhoB sensor (Supplementary Fig. 1a (iii),b), indicating that the activities reported by RhoA-based probes were affected by local cell geometry. Thus, ratiometric FRET analysis based on epifluorescence imaging data insufficiently corrects for cell geometry when most of the inactive probe resides in the cytoplasm, as in the case of RhoA-based probes. To further explore the suitability of the RhoB sensor for spatiotemporal analysis of Rho activity, we tested its response to plasma membrane translocation of mCherry-FKBP-ARHGEF1(GEF) and mCherry-FKBP-ARHGAP29(GAP), through rapamycin-induced heterodimerization with plasma membrane-localized Lyn_11_-FRB, to acutely (in)activate Rho activity in cells (Fig. 1g,h,i). ARHGEF1 and ARHGAP29 are a Rho-specific GEF and GAP, respectively, and the RhoB sensor showed bidirectional, expected responses to plasma membrane translocation of their catalytic domains. Thus, due to the superior response to thrombin, the absence of a noticeable localization bias, and the robust responses to both activating and inactivating Rho-regulators, we considered the RhoB sensor as the preferred Rho probe for spatiotemporal analysis in HT-HUVEC and our imaging systems for subsequent analysis.

As Rac and Cdc42 reporters, we chose RaichuEV-Rac and RaichuEV-Cdc42^19,42,48,49^. They reported expected responses to pathway-stimulating signals. Increased activities occurred upon plasma membrane translocation of mCherry-FKBP-TIAM(GEF) (Fig. 1j) and mCherry-FKBP-ITSN1(GEF) (Fig. 1k) with TIAM being a Rac-specific and ITSN1 a Cdc42-specific RhoGEF. We noted that RaichuEV-Rac and RaichuEV-Cdc42 co-activated in response to the GEF translocations, consistent with positive feedback mechanisms that engage during cell protrusions^50^. RaichuEV-Rac and RaichuEV-Cdc42 were inactivated and activated upon Rho activation and inactivation, respectively (Fig. 1f,i) (with the exception for ARHGEF1-activated Rho, Fig. 1h), consistent with an antagonism between protrusive and contractile RhoGTPase activities^51^. In addition to expected bidirectional responses of all RhoGTPase probes tested, their temporal responses were on the order of seconds, further validating their suitability for spatiotemporal analysis in motile cells.

We then imaged randomly motile HT-HUVEC stably expressing RaichuEV-Rac, RaichuEV-Cdc42, RhoB sensor, DORA-RhoA, or RhoA2G or at 25s intervals to identify spatiotemporal activation patterns of Rho GTPases during protrusion-retraction cycles (Fig. 2, Videos 2-6). For each probe, panels (i) show RhoGTPase activity differences between thresholded protrusions and retractions, normalized to non-moving membrane segments in the same time period. Panels (ii) show single cell spatiotemporal maps visualizing edge velocities and RhoGTPase activities within 1.95µm from the cell edge, and cross-correlation between the two maps. Panels (iii) show cross correlations between edge velocity and RhoGTPase activities compiled from multiple cells. As expected, relative to membrane segments classified as non-moving, Rac1 and Cdc42 were elevated in protrusions and reduced in local retractions (Fig. 2a,b (i)). Cross-correlation analysis between edge velocity and Rac1 (Fig. 2a (iii)) or Cdc42 (Fig. 2b (iii)) showed strong positive correlations with lags of negative 25-50s (i.e. peak RhoGTPase activities lagging peak edge velocities), confirming previous results^14,15^. Interestingly, the RhoB sensor showed an opposite pattern of activity - uniformly inactive in protrusions, and variably but consistently activated in retractions (Fig. 2c (i)). Cross-correlation between edge velocity and RhoB sensor activity yielded a sharply negative peak lagging edge velocity by 25-50s (Fig. 2c (iii)). Similar analyses using DORA-RhoA or RhoA2G confirmed enrichment of Rho activity in retractions (Fig. 2d,e (i)). Cross-correlations between edge velocity and RhoA2G or DORA-RhoA signals were less uniform between cells, but similarly consistently negative (Fig. 2d,e (iii)), confirming that edge-proximal Rho activity is correlated with local edge retractions and anticorrelated with protrusions. Together, our analyses demonstrate that µm and minute-scale cell shape changes in randomly motile HT-HUVEC are associated with distinct RhoGTPase activity patterns, with elevated Rac and Cdc42 activities protrusions and elevated Rho activity in retractions.

**Figure 2.**
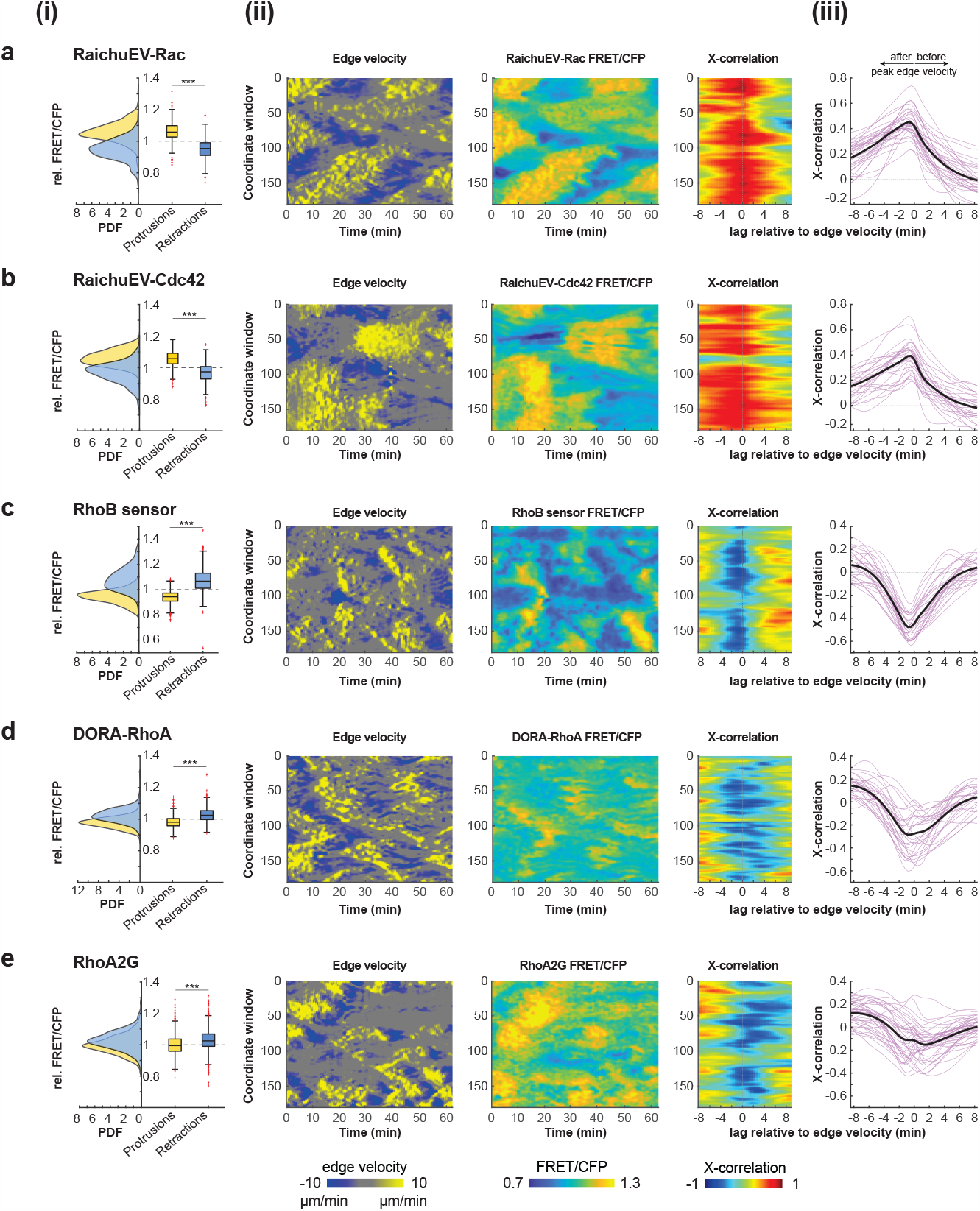
Spatiotemporal analysis of Rac, Cdc42, and Rho activities in HT-HUVEC. (**a)** RaichuEV-Rac. **(b)** RaichuEV-Cdc42. **(c)** RhoB sensor. **(d)** DORA-RhoA. **(e)** RhoA2G (Videos 2-6). **(i)** Graphs showing differences in relative FRET activity in thresholded protrusions and retractions for each probe tested at a window edge depth of 1.95µm. Protrusions > 3.86µm/min, retractions < -3.86µm/min, each minimum 25 pixels on edge velocity heat maps. Individual protrusions and retractions were normalized to edge-proximal FRET activity in non-moving edge areas during the same time span. Left x axis shows probability density functions of retractions in blue, and protrusions in yellow. Box plot on the right x-axis: bolded center line denotes dataset median, coloured boxes show 25th and 75th percentiles, and whiskers show total dataset range. Outliers are shown with red crosses. Data for each probe were from 3 biological replicates. RaichuEV-Rac: 30 cells, 489 protrusions, 299 retractions. RaichuEV-Cdc42: 29 cells, 552 protrusions, 308 retractions, RhoB sensor: 36 cells, 726 protrusions, 413 retractions. DORA-RhoA: 30 cells, 771 protrusions, 672 retractions, RhoA2G: 40 cells, 1100 protrusions, 1021 retractions. *\*\*\*p<0*.*001*, Mann-Whitney U-test. **(ii)** Examples of spatiotemporal heat maps for individual cells, from 62.5min time-lapse acquisitions. (Left) edge velocity, with protrusions in yellow and retractions in blue, and (center) corresponding RhoGTPase activity, measured within 1.95µm from the cell edge. (Right) cross-correlation between edge velocity and RhoGTPase activity from the two heatmaps. **(iii)** Cross-correlation of RhoGTPase activity relative to edge velocity for each probe at a window depth of 1.95µm. Pink traces represent single cells, bold black traces are means. Negative lag denotes peak RhoGTPase activity following peak edge velocity and vice versa. RaichuEV-Rac n=34 cells, RaichuEV-Cdc42 n=28 cells, RhoB sensor n =27 cells, from 2 biological replicates. DORA-RhoA n=29 cells, RhoA2G n=40 cells, from 3 biological replicates.

### Pulsatile activation of Rho during membrane retractions

We subsequently focused on Rho activity in membrane retractions, since Rac and Cdc42 have well-documented roles in driving membrane protrusions. Visually, RhoB sensor activity propagated from the cell interior towards the cell edge before retraction onset (Fig. 3a,b, Video 7). Our cross-correlation analyses between edge velocities and Rho activity yielded a negative correlation and time-lag, however, these parameters result from averaged relationships in protrusions and retractions combined. To specifically determine the kinetics of Rho activation relative to membrane retractions (but not Rho inactivation in protrusions), we analyzed RhoB sensor activity in parts of spatiotemporal edge velocity maps where protrusions transitioned into retractions. (Supplementary Fig. 2, Materials and Methods). To compare RhoB sensor activation across multiple retraction events, we aligned Rho kinetics and edge velocity time courses by setting time = 0 to the transition point from protrusion to retraction (edge velocity = 0). The resulting averaged Rho activity time-course showed a striking, pulse-like activation during retractions (Fig. 3c). RhoB sensor activity began to rise well before retraction onset, while edge velocity remained positive. Upon reaching the transition point, RhoB sensor levels were already well above basal activation levels and increased in a non-linear fashion. Surprisingly, RhoB sensor activity did not remain statically elevated throughout the retraction. Peak RhoB sensor activity 2 min after retraction onset was followed by a marked reduction and return to baseline levels, indicating the presence of an inactivation mechanism triggered before retraction completion.

**Figure 3.**
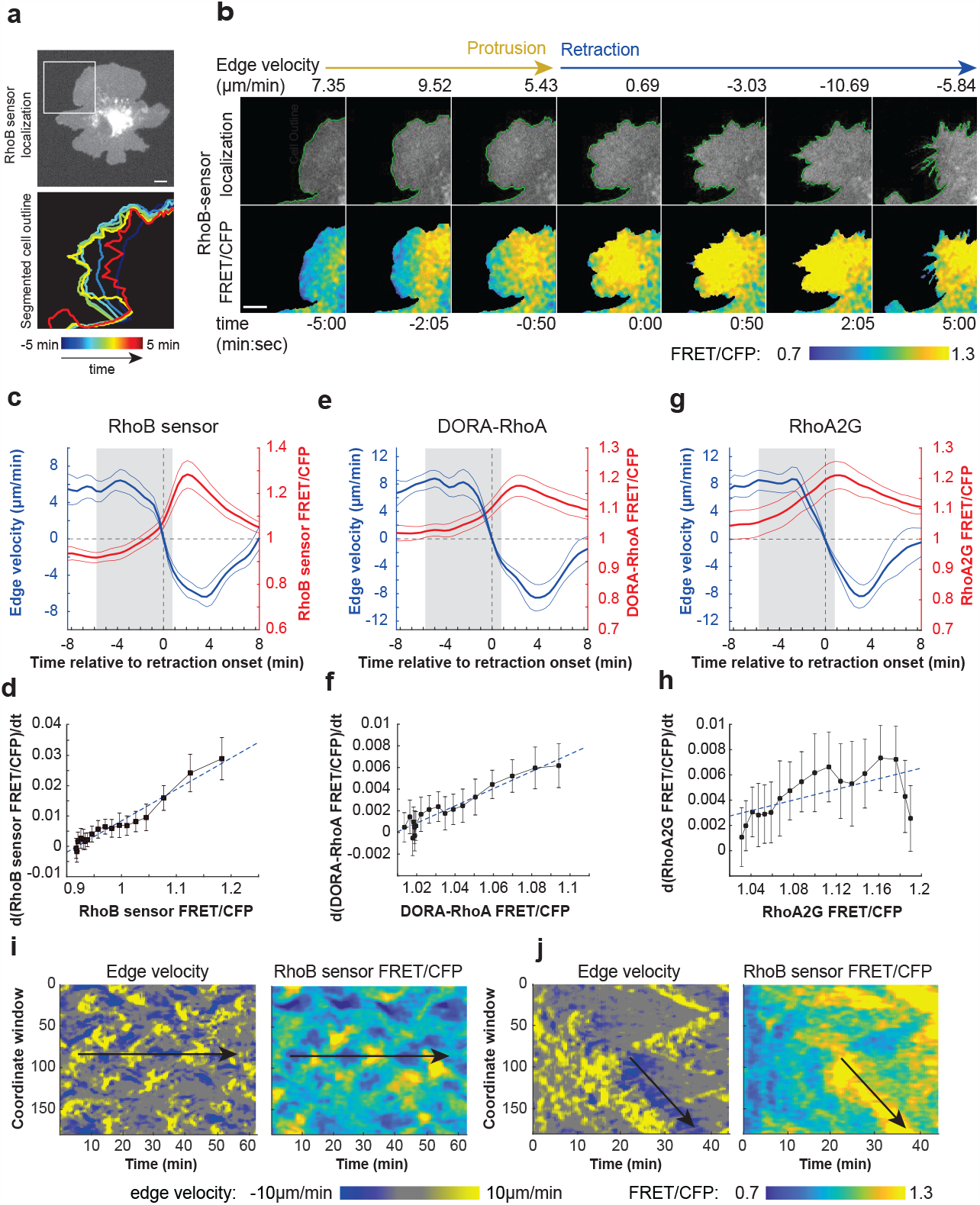
Pulsatile activation of Rho in cell edge retractions. (**a)** *(Top)* HT-HUVEC, expressing the RhoB-sensor, with region of interest denoted by white box. Scale bar, 10µm. *(Bottom)* local cell-edge displacements over 10min of the protrusion retraction event shown in (b). **(b)** Time-lapse illustrating RhoB sensor activation during a protrusion retraction event during a 10min interval. Time=00:00 denotes retraction onset. Scale bar, 10µm. (Video 7). **(c, e, g)** Activity buildup plots displaying averaged RhoB sensor, DORA-RhoA, and RhoA2G activities, respectively, compared during protrusion-retraction transitions. Time = 0 denotes the protrusion-retraction transition. Grey shaded region shows timepoints included in the rate-of-change analysis in d,f,h. Means ± 95 CI. RhoB sensor: n=23 events, DORA-RhoA: n=33 events, RhoA2G: n=25 events, from 3 biological replicates. **(d, f, h)** Plot of the rate-of-change in Rho activation as a function of Rho activity from t= -7min to t = 1.5min (18 25s timepoints) from the buildup plots for RhoB sensor, DORA-RhoA, and RhoA2G. Black boxes denote timepoint means, error bars show ± 95% CI. Linear line of best fit shown by dotted blue line. **(i, j)** *(Left)* Thresholded edge velocity and *(Right)* RhoB sensor activity maps (window depth 1.95µm) of representative cells illustrating (i) pulsatile activity and (j) a propagating wave of RhoB sensor activity. Region of interest denoted by black arrows on both sets of maps (Videos 8, 9).

Such pulse-like patterns of activity are generated by activator-inhibitor coupled excitable systems, comprised of fast auto-activation paired with delayed self-inhibition^52–54^. Plotting the rate of change of RhoB sensor activity as a function of RhoB sensor activity during the buildup phase (approximately 6min before retraction onset to 1min post retraction onset, shaded rectangle in Fig. 3c) revealed a linear positive correlation (Fig. 3d), i.e., Rho activity increase accelerating with increasing Rho activity. This is consistent with active Rho engaging in a positive feedback loop to further increase its own activation, a phenomenon noted in other instances of excitable Rho behavior^55,56^. In addition, we observed pulsatile behavior (Fig. 3i, Video 8) and propagating waves (Fig. 3j, Video 9) of both RhoB sensor activity and edge retraction, patterns consistent with an excitable Rho signalling network. To confirm this finding, we repeated Rho buildup analyses using the DORA-RhoA and RhoA2G probes and observed similar pulsatile activation patterns and accelerating positive feedback phases, albeit on a smaller scale (Fig. 3e-h). This was likely due to RhoA2G and DORA-RhoA ‘s reduced dynamic range, and cell geometric bias artificially increasing Rho activity near the cell edge (Supplementary Fig. 1). Buildup profiles of Rac and Cdc42 activity in protrusion-retraction transitions showed a decrease in activity aligned with retraction onset but no pulsatile patterning (Supplementary Fig. 3). Thus, Rho activity increases before retraction onset and is pulsatile, with rapid activation during and inactivation before the retraction process is completed.

### Rho, ROCK, NMII are part of an excitable system during membrane retractions

Biological excitability is characterized by positive self-feedback which generates a rapid increase in activity, followed by delayed inhibition by a downstream effector that returns activity to basal levels^53,54^. Excitable signalling modules can drive cell protrusions during chemotaxis or propagating waves of actin polymerization^19,53,57–59^. Moreover, previous studies have found that Rho and its cytoskeletal effectors are excitable, resulting in pulsatile or wave-like propagation of Rho activity^55,56,60,61^. Interestingly, several studies identified that accumulation of the downstream target NMII with a delay relative to Rho can act as a negative regulator of Rho through recruitment of RhoGAPs^55,56,60–62^. To test whether this was the case during membrane retractions, we first coexpressed fluorescently tagged myosin regulatory light chain (mRuby3-MLC) and RhoB sensor in HT-HUVEC to analyze myosin activation alongside edge velocity and Rho activity. The localization-based myosin activity reporter mRuby3-MLC is cytoplasmic when inactive but forms distinct puncta when incorporated into myosin filaments upon activation. Spatiotemporal analyses revealed that edge-proximal mRuby3-MLC signal was low during protrusions and early retractions (Fig. 4a, Video 10). Levels increased in intensity beginning at retraction onset, peaking several minutes after maximal Rho activity levels (Fig. 4b). Cross-correlation analysis between RhoB sensor and MLC-mRuby3 signal showed a delayed positive correlation (Supplementary Fig. 4a), significantly highest at edge depths of 5-8µm (Supplementary Fig. 4b,c), reflecting the absence of edge-proximal myosin and localization to inward moving actin filaments. The slow, delayed accumulation of myosin following Rho activation in protrusion-retraction events closely matched the localization of negative regulators in excitable system models^53^, consistent with NMII activation downstream of Rho activity being part a negative regulator of Rho itself.

**Figure 4.**
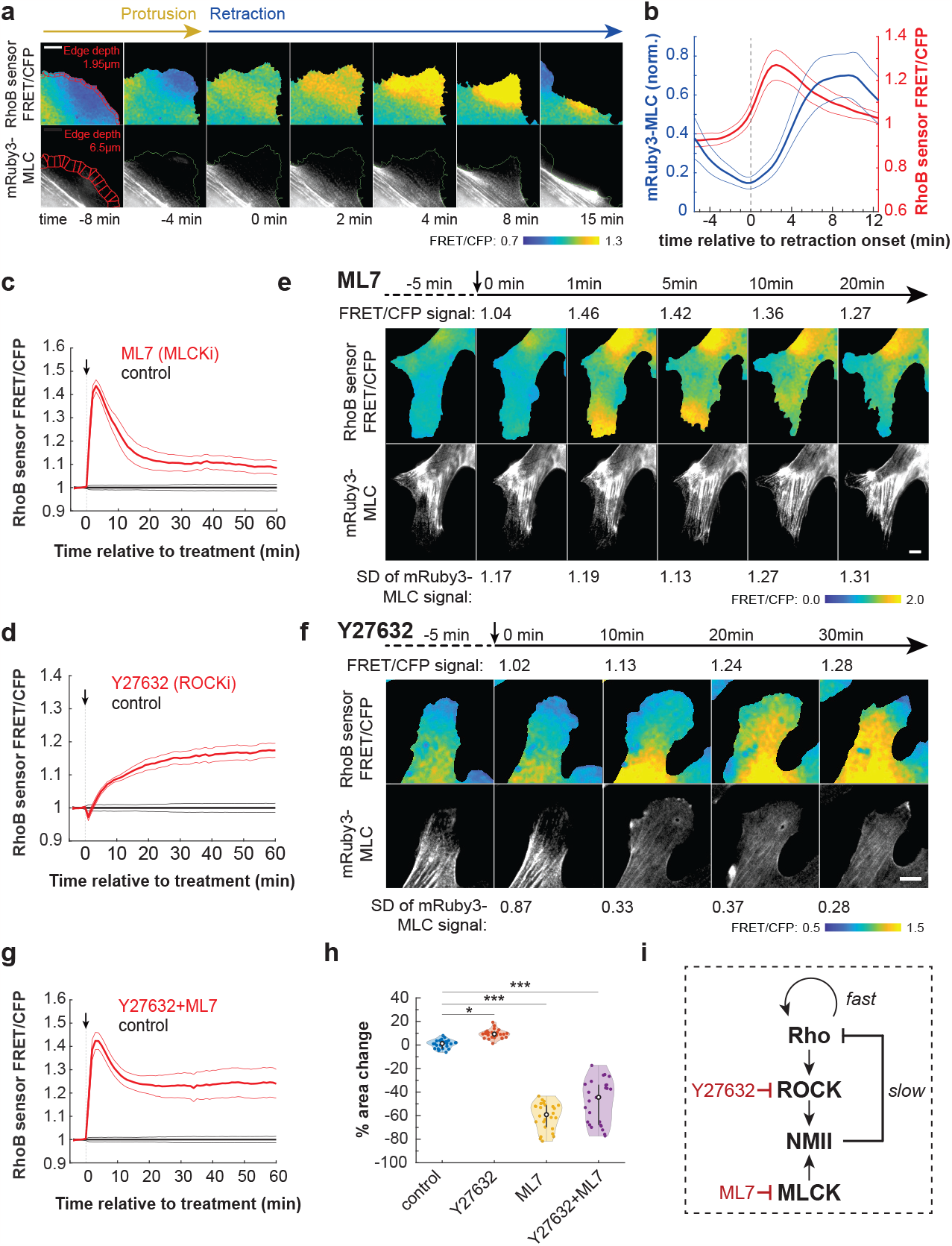
Rho, ROCK, NMII are part of an excitable system during membrane retractions. (**a)** Time-lapse showing Rho activation and NMII accumulation in a RhoB-sensor and mRuby3-MLC-expressing HT-HUVEC. Red edge windows show 1.95µm window depth for RhoB sensor and 6.5µm for MLC-mRuby3. t=0 denotes retraction onset. Scale bar, 10µm. (Video 10). **(b)** 30 protrusion-retraction events, from 3 biological replicates. RhoB sensor window depth, 1.95µm, mRuby3-MLC window depth, 6.50µm. Bolded lines represent means, bordered by ± 95% CI. **(c**,**d**,**g)** RhoB sensor activity responses to acute inhibition of MLCK (ML7, 20µM), ROCK (Y27632, 20µM), or ROCK and MLCK (Y27632+ML7, both 20µM). Drug responses were normalized to control-treated samples. Bolded lines denote means, bordered by ± 95% CI. 26 fields of view (FOV) for each condition, from 3 biological replicates. **(e)** Effect of MLCK inhibition (ML7, 20µM) on RhoB sensor activity *(top)* and mRuby3-MLC distribution *(bottom)*. Rho signal labelled above top row, and standard deviation of pixel distribution to measure granularity of MLC signal labelled below bottom row. Scale bar, 10 µm. (Video 11). **(f)** Effect of ROCK inhibition (Y27632, 20µM) on RhoB sensor activity *(top)* and mRuby3-MLC distribution *(bottom)*. RhoB sensor activity labelled above top row, and standard deviation of pixel distribution to measure granularity of mRuby3-MLC signal labelled below bottom row. Scale bar, 10µm. (Video 12). **(h)** Percent area change of cell adhesion surface 30min post addition of either control, 20µM Y27632, 20µm ML7, or a combination of 20µM Y27632 and 20µM ML7. n=26 FOV each for control, Y27632, and ML7 conditions, from 3 biological replicates. ^*\**^*p<0*.*05*, ^*\*\*\**^*p<0*.*001*, one-way ANOVA/Tukey-Cramer. **(i)** Diagram illustrating the relationship between Rho ‘s downstream effectors ROCK and NMII and the drugs used in c-h.

To test this further, we measured RhoB sensor activity in response to acute treatments of cells with inhibitors of ROCK (Y27632, 20µM) and myosin light chain kinase (MLCK, ML7, 20µM), i.e., kinases acting upstream of NMII activation. We were unable to use blebbistatin as a direct inhibitor of myosin motor activity, as the color of solutions containing blebbistatin interfered with FRET-based Rho activity measurements. Consistent with NMII acting as a negative regulator of Rho, both drug additions resulted in increased RhoB sensor activity, although with distinct kinetics. ML7 addition resulted in a sharp, transient increase (Fig. 4c), whereas Y27632 addition caused a slower, but persistent increase of Rho activity (Fig. 4d). Within MLCK-inhibited cells, stress fibers were maintained or increased, and RhoB sensor activity transiently increased in peripheral regions (Fig. 4e, Video 11), whereas ROCK inhibition caused sustained accumulation of RhoB sensor activity throughout the cells, particularly where stress fibers had dissolved, and except for peripheral ruffling regions (Fig. 4f, Video 12). Treating cells with both drugs resulted in a combined response reflective of the two independent pathways of MLC phosphorylation (Fig. 4g). Consistent with ROCK and MLCK preferentially activating NMII in the cell interior and the cell periphery, respectively, ROCK inhibition increased and MLCK inhibition decreased the cells’ adhesion surface^63,64^ (Fig. 4h). Together, both inhibitors increased Rho activity, demonstrating generally that activated NMII negatively regulates Rho. Specifically, the sustained increase in Rho activity following ROCK inhibition and stress fiber dissolution suggested that the Rho-ROCK-NMII pathway plays a critical role as a negative regulator of excitable Rho activity (Fig. 4i).

### Rho activity and ERM activation are spatiotemporally coupled

Given that mRuby3-MLC signal was at its lowest near the cell edge at retraction onset and accumulated with a substantial delay relative to Rho (Fig. 4b), we questioned whether additional Rho effectors besides NMII were involved in driving early phases of membrane retractions. Rho is also known to activate ERMs through the kinases SLK and LOK^34,35,38^. Since release of cortical actin from the plasma membrane is necessary for the initiation of membrane protrusions^42,43^, we hypothesized that re-establishment of membrane-cortex attachment through activation of ERMs could be involved in retraction initiation. To test this, we first performed live-cell imaging of RhoB sensor and mRuby3-MLC-expressing HT-HUVEC, then fixed the cells and stained them using anti-pERM antibody that detects activated (phosphorylated) ERMs. Matching membrane retractions from time-lapse sequences to pERM signals revealed a striking enrichment of pERM in retractions, consistent with Rho locally activating ERMs (Fig. 5a, Video 13). We also found that in ROCK and MLCK inhibited cells, locally increased Rho activity colocalized with locally increased pERM signals, further supporting Rho activity being upstream of ERM activation (Fig. 5b).

**Figure 5.**
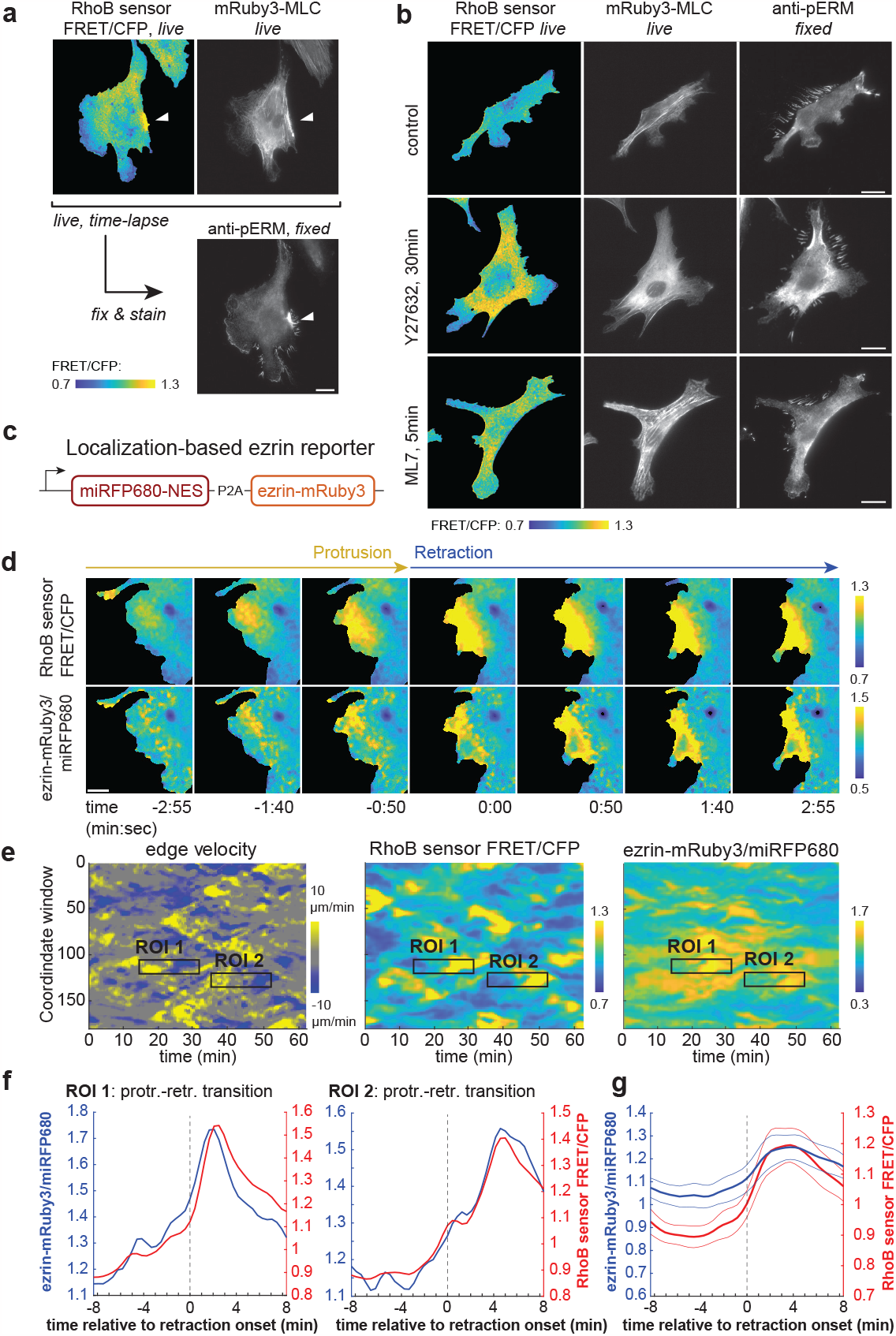
Rho activity and ERM activation are spatiotemporally coupled. (**a)** HT-HUVEC co-expressing RhoB sensor and mRuby3-MLC showing RhoB FRET activity *(live)*, mRuby3-MLC *(live)*, and corresponding anti-pERM signals following fixation and immunostaining. White arrowhead denotes retraction site. Scale bar, 20µM. (Video 13). **(b)** HT-HUVEC co-expressing RhoB sensor and MLC-mRuby3 were imaged either after no treatment, 30min post 10µM Y27632 addition, or 5min post 5µM ML7 addition, fixed, and processed for anti-pERM immunostaining. Drug-induced RhoB sensor activity increases colocalized with increased pERM staining. Scale bars, 20µm. **(c)** Schematic of the localization-based ezrin reporter. **(d)** Time-lapse showing RhoB sensor activity and ezrin-mRu-by3/miRFP680 ratio during an individual protrusion-retraction transition. Time=0:00 denotes retraction onset. Scale bar, 10µm. (Video 14). **(e)** Spatiotemporal heat maps depicting edge velocity, RhoB sensor activity, and ezrin-mRuby3/miRFP680 ratio, both measured at 1.95µm window depths. Region of interest (ROI) 1 and 2 correspond to the regions used to generate the correspondingly labelled individual activity buildup plots in (f). **(f)** Protrusion-retraction events outlined in (e) displaying average ezrin-mRuby3/miRFP680 ratios in blue and averaged RhoB sensor activities in red. Time=0 denotes retraction onset. **(g)** Compiled protrusion-retraction transitions showing normalized ezrin-mRuby3/miRFP680 ratios in blue and RhoB sensor activities in red. Time=0 denotes protrusion-retraction transition. Bolded lines represent means, bordered by ± 95% CI. 19 events from 3 biological replicates.

We next sought to monitor the spatiotemporal relationship between Rho and ERM activation during membrane retractions in live cells. We generated a localization-based ERM activation reporter, composed of mRuby3-tagged ezrin and cytoplasmic miRFP680 that we expressed stoichiometrically from the same mRNA through the use of a P2A ribosomal skipping sequence^65^ (Fig. 5c). Since ERMs are recruited to the plasma membrane upon activation^39^, the pixel-by-pixel ratio of ezrinmRuby3/miRFP680 fluorescence is an indication of local ezrin activation. Analysis of cell-edge dynamics, RhoB sensor activity, and ezrin-mRuby3/miRFP680 together revealed a striking spatiotemporal correlation of Rho activity and ezrinmRuby3/miRFP680 (Fig. 5d,e, Video 14). Activity buildup plots of both individual and compiled (Fig. 5f,g) protrusion-retraction events demonstrated no detectable time lag between Rho activation and ezrin signal in retractions. This tight spatiotemporal correlation between Rho activity and active ezrin suggested that ERMs are immediate-early effectors of Rho, which serve to enhance the force transmission between actin and the plasma membrane that is necessary during retractions.

### Rho rapidly activates ERMs via SLK/LOK

To test whether SLK/LOK kinases are involved in ERM activation downstream of Rho in HT-HUVEC (Fig. 6a), we first characterized a recently developed SLK/LOK inhibitor (Cpd31)^66^. Treatment of HT-HUVEC with Cpd31 caused an almost complete loss of pERM signal, as assessed by quantitative immunofluorescence, with only some detectable pERM signal remaining in peripheral structures resembling retraction fibers (Fig. 6b). No major changes in the actin cytoskeleton of treated cells were apparent (Fig. 6b). pERM signal decreased dose-dependently (Fig. 6c,d), and occurred within 2.5min of Cpd31 addition at 5µM (Fig. 6e,f). This surprisingly rapid loss of pERM upon SLK/LOK inhibition indicated a high turnover rate of ERM phosphorylation^38^. We next used acute treatment of cells with thrombin or nocodazole to assess Rho-dependent changes in pERM signals (Fig. 6a). Thrombin and nocodazole both rapidly activate Rho: Thrombin through the GPCR PAR-1^67^; nocodazole through release of microtubule-sequestered GEF-H1, a Rho-specific GEF, from depolymerizing microtubules^68,69^. Both treatments caused rapid, concurrent increases in RhoB sensor activity (Fig. 1e, Supplementary Fig. 5a) and pERM signals (Fig. 6g,h). Strikingly, the increases in pERM signal were potently suppressed when Cpd31 was present (Fig. 6h,i). Treating cells with ROCK inhibitor (Y27632) caused no significant reduction in pERM signals, indicating that ROCK is not required for ERM activation in HT-HUVEC (Fig. 6g,h). Our results therefore demonstrate an essential role of SLK/LOK in rapid ERM activation downstream of Rho and argue for the Rho-SLK/LOK-ERM signaling axis being a highly responsive signaling module for the regulation of actin-membrane attachment and force transmission.

**Figure 6.**
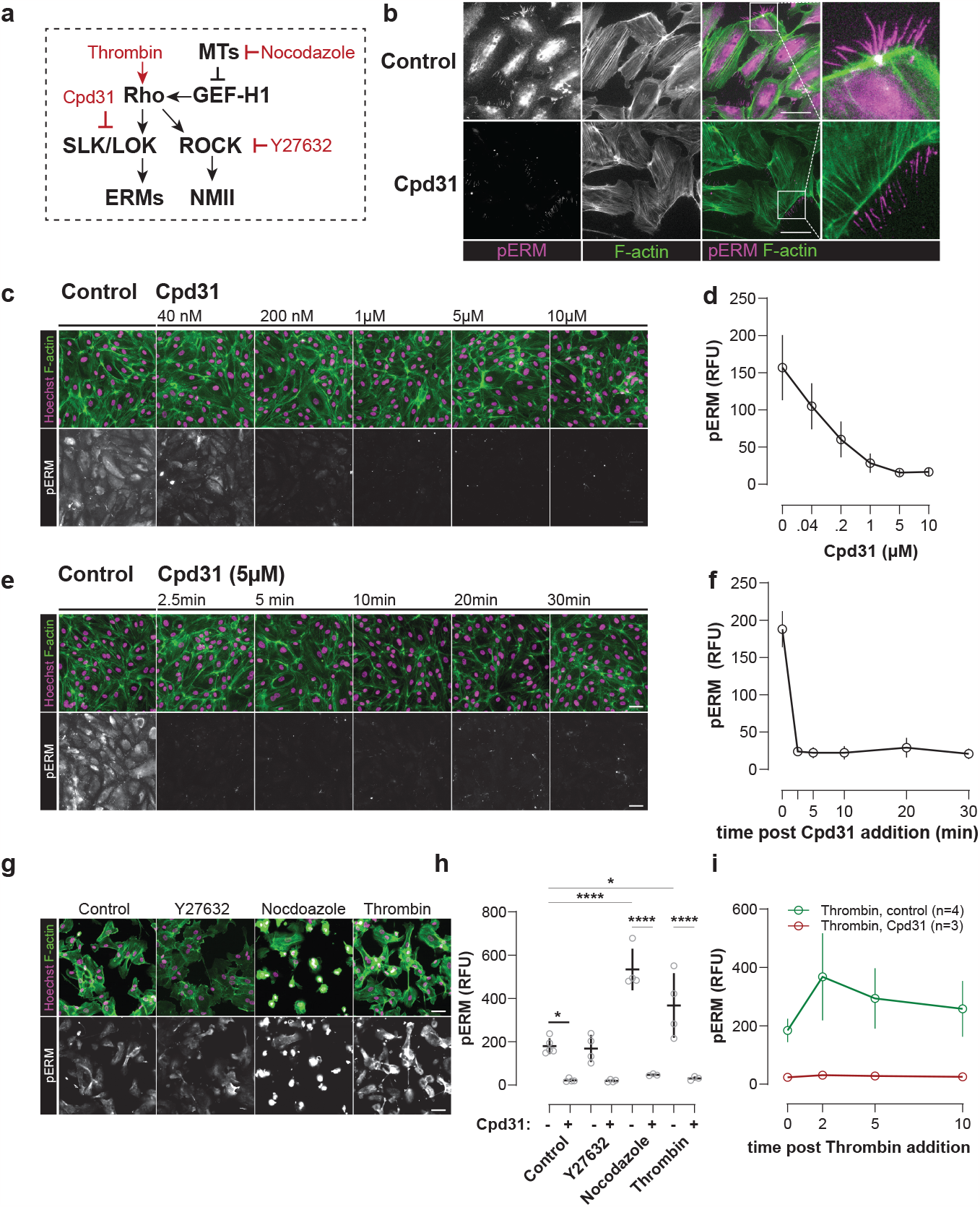
Rho rapidly activates ERMs via SLK/LOK. (**a)** Perturbation strategies to investigate Rho-induced ERM activation. **(b)** HT-HUVEC treated with the SLK/LOK inhibitor Cpd31 (5µM, 4h) show an almost complete loss of pERM signal, whereas F-actin organization appears unaffected. **(c)** pERM signal in HT-HUVEC in response to increasing concentrations of Cpd31 (4h). **(d)** Quantification of (c) using quantitative immunofluorescence (Materials & Methods), means ± SD from n=3 biological replicates. **(e)** Cells rapidly lose pERM signals when treated with Cpd31 (5µM). **(f)** Quantification of (e) using quantitative immunofluorescence, means ± SD from n=3 biological replicates. **(g)** Treatment with nocodazole (15µM, 30min) or thrombin (1U/ml, 2min) increased pERM in HT-HUVEC, whereas Y27632 (10µM, 30min) had no detectable effect. **(h)** Nocodazole and thrombin-induced increases in pERM were abolished when Cpd31 (5µm) was present. Means ± SD from n=3-5 biological replicates are shown. ^*\**^*p<0*.*05*, ^*\*\*\*\**^*p<0*.*0001*, one-way ANOVA/Tukey-Cramer. **(i)** pERM signals rapidly increased upon thrombin stimulation (1U/ml), and pERM increases were abolished when Cpd31 (5µM) was present. Means ± SD from n=4 (thrombin) or n=3 (thrombin+Cpd31) biological replicates. Scale bars, 50µm.

### SLK/LOK dependent ERM activation regulates cell morphology and is required for Rho-driven cell contractions

Given the importance of Rho activity for cell shape homeostasis and cell contractions, we took advantage of the ability to selectively inhibit the Rho downstream targets ROCK/NMII (using Y27632) and SLK/LOK/ERM (using Cpd31) to identify their individual or combined roles in these processes.

For cell shape homeostasis, we plated HT-HUVEC without, or in presence of Cpd31, Y27632, or Cpd31 and Y27632 combined, fixed the cells after 2h, and stained them with the AF647-conjugated surface label wheat germ agglutinin (WGA647) (Fig. 7a). The advantage of this protocol is that cells have not yet assumed heterogenous cell morphologies prior to exposure to the inhibitors, facilitating the detection of morphological phenotypes. Importantly, for all of the conditions tested, cells attached and spread to similar adhesion areas (Fig. 7c). Y27632 treatment is known to cause elongated cell morphologies and retraction defects^70–72^.We found that inhibition of either ROCK or SLK/LOK had similar effects, with cells having extended cellular processes (Fig. 7a). To quantify drug-induced cell shape changes, we chose eccentricity and solidity as metrics. Eccentricity quantifies cell elongation and solidity is the ratio of cell area and the area of a cell ‘s convex hull (cell compactness) (Fig. 7b, Materials and Methods). Both Cpd31 and Y27632 yielded significantly increased eccentricity and significantly reduced solidity (Fig. 7d,e), and combined treatment had additive effects on both metrics (Fig. 7d,e). These results indicated that the Rho effectors ROCK and SLK/LOK both contribute to cell shape homeostasis by restricting extended cellular processes.

**Figure 7.**
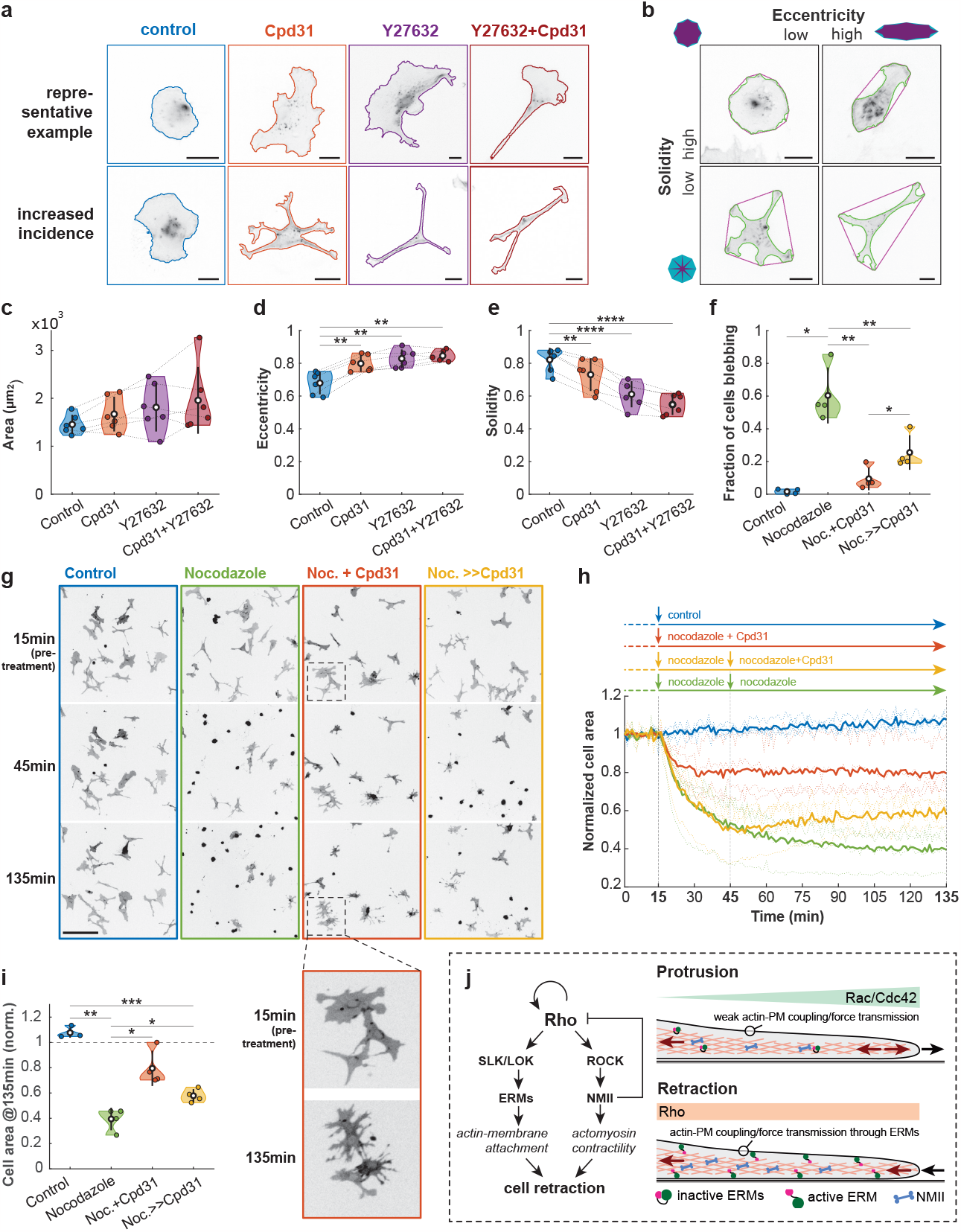
SLK/LOK-dependent ERM activation regulates cell morphology and is required for Rho-driven cell contractions. **(a)** HT-HUVEC were plated without (control) or in presence of inhibitors of SLK/LOK (Cpd31, 5µM), ROCK (Y27632, 10µM) or SLK/LOK & Y27632 combined, fixed after 2h and stained with WGA-AF647. *(Top)* Representative examples of cell shapes with average morphologies and *(bottom)* examples of cell shapes frequently observed in the presence of the inhibitors are shown. Scale bars, 20µm. **(b)** Illustrations of cell shapes with low and high solidity and eccentricity. Scale bars, 20µm. **(c)** Average cell spreading area, **(d)** eccentricity, and **(e)** solidity. Data points are means from n=6 biological replicates (control: 1141 cells, Cpd31: 966 cells, Y27632: 776 cells, and Cpd31 & Y27632: 696 cells total). ^*\*\**^*p<0*.*01*, ^*\*\*\*\**^*p<0*.*0001*, one-way ANOVA/Tukey-Cramer. **(f)** Fraction of HT-HUVEC expressing RhoB sensor blebbing after either control treatment, 2h nocodazole (15µM), 2h nocodazole (15µM) & Cpd31 (5µM), or 2h nocodazole (15µM), with Cpd31 (5µM) added after 30min (as depicted in schematic in (h), t=135 min). (g) HT-HUVEC expressing RhoB sensor were treated as depicted in the schematic in (h). Nocodazole (15µM), Cpd31 (5µM). RhoB sensor localization was used to visualize cells. Zoom-in shows fragmented cells in nocodazole & Cpd31 condition. Scale bar, 200µm. (Video 15). **(h)** Average area covered by cells per field of view (FOV) as a function of time, normalized to the average cell area of the first 14min for each condition. First treatment at 15min, second treatment at 45min (black dashed lines). Each bold trace represents the means of n=4 biological replicates (replicate means shown as colored dashed lines). **(i)** Average cell-covered area per field of view in h at time 135min for each condition, normalized to pre-treatment. (f,h,i) Means from n=4 biological replicates (control: 14 FOV, nocodazole: 13 FOV, nocodazole + Cpd31: 14 FOV, and nocodazole followed by Cpd31: 14 FOV analyzed in total). ^*\**^*p<0*.*05*, ^*\*\**^*p<0*.*01*, ^*\*\*\**^*p<0*.*001*, one-way ANOVA/Tukey-Cramer. **(j)** Model for how excitable Rho activity drives cell contractions through both ERM activation via SLK/LOK and ROCK-mediated actomyosin contractility *(see main text)*.

To directly assess the role of Rho-driven ERM activation in membrane retractions, we treated fully adhered cells with nocodazole, as it activates Rho and causes a reduced adhesion surface of cells (Supplementary Fig. 5a,b). Upon nocodazole addition, cells rapidly contracted and many started blebbing (Fig. 7f,g, Video 15). Strikingly, when Cpd31 was added along with nocodazole, both the fraction of cells blebbing and cell contractions were significantly reduced, demonstrating that SLK/LOK are required for cells to exhibit a contractile response downstream of Rho (Fig. 7f,g,h,i). In samples where cells were first treated with nocodazole for 30min to induce Rho-dependent blebbing, the addition of Cpd31 increased cell spreading and a significant fraction of cells recovered from blebbing (Fig. 7f,g,h,i, Video 15). This suggested that the process of bleb retraction not only resolves blebs, but also maintains blebbing cells in a blebbing state. Intriguingly, cells treated with combined nocodazole and Cpd31 often had “runaway” protrusions resulting in cell fragments that detached from the main cell body (Fig. 7g). Importantly, Cpd31 treatment did not prevent nocodazole-induced Rho activation (Supplementary Fig. 5c).

Together, our results demonstrate that the Rho effectors ROCK and SLK/LOK are both critical for Rho-induced cell contractions. Moreover, SLK/LOK activation of ERM downstream of Rho appears to be critical to safeguard cell integrity.

## Discussion

By analyzing the spatiotemporal dynamics of Rho activity in randomly motile HT-HUVEC, we found that Rho was consistently elevated in µm and minute-scale cell edge retractions and consistently absent from protrusions. That Rho was elevated in retractions was confirmed by the use of three different FRET probes and is consistent with both recent studies using FRET-based and localization-based reporters for Rho^21,22^ and the generally accepted role of Rho in activating NMII-dependent cell contractions through ROCK.

Intriguingly, our spatiotemporal analysis revealed that Rho activity in retraction events displays hallmark characteristics of excitability, consistent with fast positive and slow/delayed negative inhibition. This finding is supported by the multiple instances of excitable Rho activity generating contractile pulses in non-migratory contexts^55,56,60,61^.

By including MLC or the ERM protein ezrin in our spatiotemporal analyses of Rho dynamics, we found that MLC accumulated with a minute-scale delay at retracting cell edges relative to Rho activity, whereas ezrin showed high co-localization without detectable delay. In previous studies on excitable Rho dynamics, actomyosin-associated RhoGAPs have been identified as negative regulators of Rho, in line with our findings that myosin inhibition increased Rho activity^56,60,73^. Therefore, it is likely that an actomyosin-associated RhoGAP is recruited to retractions as they evolve, progressively shutting off Rho and ensuring contractions stop.

While positive feedback to Rho most likely involves the recruitment of a RhoGEF^55,60,61,73,74^, it is less clear how this occurs during retractions. The tight spatiotemporal coupling between Rho and ezrin demonstrates that ERMs are early Rho effectors during retractions and indicates they could be components of a positive feedback mechanism. Both positive and negative feedback from ERMs to Rho activity have been observed^35,75,76^. However, negative feedback through ERMs is unlikely in our case, given the absence of delay between Rho and ezrin activation. Focal adhesion dynamics may also contribute to regulating Rho during protrusion-retraction cycles. Focal adhesions have been shown to recruit RhoGEFs and GAPs in a maturation-dependent manner, with RacGEFs associated with newly formed adhesions near the cell edge and RacGAPs and RhoGEFs with mature adhesion sites deeper in the cell interior^77^. Furthermore, increased tension on focal adhesions can recruit the RhoGEFs LARG and GEF-H1^78^, meaning greater Rho-induced contractility leads to higher Rho activity. Finally, local microtubule disassembly induces retractions^79^, and the interaction between focal adhesions and microtubules can control Rho activity through the sequestration and release of GEF-H1^80^, further supporting a plausible role for GEF-H1 in Rho ‘s positive feedback.

What are the specific roles of ERMs and NMII during Rho-dependent cell edge retractions? The initial absence of NMII in early phases of retractions, despite elevated Rho, can be explained by the absence of NMII filaments in branched lamellipodial actin networks^81^. Our results are consistent with a model in which ERMs are the main immediate effectors of Rho through SLK/LOK during early phases of cell edge retractions, by re-establishing a link between plasma membrane and peripheral actin networks (Fig. 7j). In adherent cells, these networks typically move radially inwards, due to NMII contractility and/or polymerization-driven treadmilling^82–84^, which in our model drags the membrane inwards once Rho-activated ERMs enable force transmission, with ERMs acting as the actin-PM clutch. Actin network remodeling occurs during early phases of edge retraction, progressively allowing for accumulation of NMII filaments^8,85^, which in turn exert contractile forces to further accelerate edge retraction. Increasingly remodeled and contractile actomyosin then recruits a putative RhoGAP, which shuts down Rho, with the observed delay, to resolve the retraction.

Rho/ROCK-dependent NMII activation is important for cell migration and inhibiting ROCK causes a tail-retraction defect^70–72^. Treating cells with the SLK/LOK kinase inhibitor induced morphological phenotypes similar to those observed during ROCK inhibition, with extended cellular processes that failed to retract. The model that both NMII and ERMs are required for cell contractions was further supported by our results showing that Rho-induced cell contractions were impaired in the presence of the SLK/LOK inhibitor. Furthermore, cells that were induced to bleb in response to acute Rho activation stopped blebbing and started spreading when SLK/LOK inhibitor was added. Cells treated this way had aberrant “runaway” protrusions that occasionally detached from cells, highlighting the critical importance of Rho-dependent ERM activation through SLK/LOK for the maintenance of cellular integrity.

In summary, our results show that in endothelial cells, Rho drives membrane retractions through two mechanisms that act sequentially during the cell edge retraction process: SLK/LOK-activated ERMs enhance actin-membrane attachment and force transmission, which enable ROCK-activated NMII to then effectively pull the membrane inwards, akin to ERMs acting as a clutch and NMII as a motor. Previous studies found that initiation of cell protrusions is preceded by a local reduction in membrane-proximal actin or ezrin, and that protrusions stalled when actin-membrane attachment was enhanced through synthetic activation of ezrin^42,43^. Therefore, in cells in which the Rho-SLK/LOK-ERM module exists, edge-proximal Rho activity appears to be incompatible with driving membrane protrusions. Since RhoA has been proposed to drive cell edge protrusions by activating mDia1^14^, it will be important to investigate whether edge-proximal RhoA and ERMs are spatiotemporally correlated in these contexts.

Our findings raise several important questions to be addressed in future research. Which are the RhoGEFs and RhoGAPs that mediate Rho excitability in cell type and context-dependent manner, and is Rho excitability a general phenomenon in motile cells? We found that Rho activity is self-limiting in randomly motile cells, with activated NMII being part of a slow negative feedback mechanism. If Rho activity is excitable during directed cell migration as well, this implies Rho activity is pulsatile rather than forming a stable rear-front gradient. Beyond cell motility and given accumulating evidence for actin-membrane attachment in regulating diverse cell morphogenetic processes, including membrane tension propagation^86^, we anticipate that our identification of ERMs as highly responsive Rho effectors will have far-reaching implications for our understanding of cell morphogenesis and migration.

## Materials and Methods

### Cell culture

HT-HUVEC, generated by stable transduction of primary HUVEC from mixed (male, female) donors with hTERT have been described^31^. They were cultured in either EGM2 (Lonza CC-3162), Endothelial Cell Growth Medium 2 (PromoCell, C-22011) or in EndoGRO VEGF (Millipore Sigma, SCME002), supplemented with 50µg/ml Hygromycin. HT-HUVEC stably expressing reporter constructs were generated by lentiviral transduction, followed by antibiotic selection (10µg/ml blasticidin, or 0.5mg/ml G418). Cells expressing multiple fluorescence-based reporters were generated by sequential lentiviral transductions and antibiotics selections. Stably transduced cells were maintained in the presence of blasticidin and/or G418 as applicable. HEK293FT cells (not authenticated), used for lentivirus production, were grown in DMEM (Thermo Fisher Scientific 11965092), supplemented with 10% FBS (Corning 35-077-CV), 5% GlutaMAX (Thermo Fisher Scientific 35050061). All cells were grown in the absence of antimicrobial agents and the absence of mycoplasma in cell cultures was routinely verified using a PCR test

### Antibodies and reagents

Rabbit anti-phospho Ezrin(T567)/Radixin(T564)/Moesin(T558) (used at 1:400) was purchased from New England Biolabs (3726S), bFGF from Cedarlane Labs (CL104-02-50UG), Thrombin from Sigma Aldrich (T4648-1KU), ML7 from Enzo (BML-EI197-0010), Y27632 form Cedarlane Labs (Y1000-1MG), Rapamycin from LC Laboratories (R-5000), Nocodazole from Cedarlane Labs (13857-5), SLK/LOK inhibitor^66^ “Cpd31” from MedChem Express (HY-132868). Phalloidin conjugated to AlexaFluor 488 was from New England Biolabs (8878S), Hoechst 33342 from Thermo Fisher Scientific (H3570), AlexaFluor-conjugated secondary antibodies from Thermo Fisher Scientific, Wheat Germ Agglutinin conjugated to AlexaFluor 647 (WGA-AF647) from Thermo Fisher Scientific (W32466), Bovine Collagen Type I from Advanced Biomatrix (5005-100ML), Paraformaldehyde solution from Fisher Scientific (50-980-487), BSA from BioShop Canada (ALB001.250), hygromycin, blasticidin, and G418 were from Invivogen (ant-hg-1, ant-bl-1, ant-gn-1, respectively).

### DNA constructs

mCherry-FKBP-GEF(TIAM1) (Addgene 85156), mCherry-FKBP-GEF(ARHGEF1) (Addgene 85152), Lyn_11_-FRB (Addgene 155228) all in pCAGEN, have been described previously ^31,42^. mCherry-FKBP-GEF(ITSN1), in pCAGEN, included mCherry, FKBP, and the DH domain of human ITSN1 (amino acids 1218-1429), which was PCR amplified from a human ORFeome clone (V5.1, clone 56297). mCherry-FKBP-GAP(ARHGAP29), in pCAGEN, included mCherry, FKBP, and the GAP domain of human ARHGAP29 (amino acids 668-900), which was PCR amplified form HT-HUVEC cDNA. pLV-RaichuEV-Rac-IRES-Blast and pLV-RaichuEV-Cdc42-IRES include a condon-diversified YPet to reduce similarity with mTurquoise at the nucleotide level to enable their lentiviral transduction. Both reporters and constructs have been described previously^42,48,49^. To generate pLV-RhoA2G-IRES-Blast, RhoA2G, kindly provided by Dr. Olivier Pertz (University of Bern, Switzerland)^44^ was PCR-amplified and inserted into pLV-EF1a-MCS-IRES-Blast (Addgene 85133) using Gibson assembly^88^. DORA-RhoA, a <u>d</u>imerization <u>o</u>ptimized reporter of <u>a</u>ctivation for RhoA^89,90^, with codon-diversified dCer3 and L9Hx3 linker, and a sensor-dead version of DORA-RhoA (PKN-L59Q) were kindly provided by Dr. Yi Wu (University of Connecticut, USA). To generate pLV-DORA-RhoA-IRES-Blast, DORA-RhoA was PCR-amplified and inserted into pLV-EF1a-MCS-IRES-Blast using Gibson assembly. The RhoB sensor used in this study has been described previously^20^. Here, pLV-RhoB sensor-IRES-Blast was generated through PCR amplification of RhoB from HT-HUVEC cDNA, DORA-RhoA minus RhoA from pLV-DORA-RhoA-IRES-Blast, and both PCR products were inserted into pLV-EF1a-MCS-IRES-Blast using Gibson assembly. A sensor-dead version of the RhoB sensor was generated similarly, by using DORA-RhoA-PKN(L59Q) as a PCR template for the DORA-RhoA minus RhoA sequence. pLV-mRuby3-MLC-IRES-Neo was created by PCR-amplifying MLC (MYL9) and mRuby3 using pLV-Ftractin-mRuby3-p2A-mTurquoise-MLC-IRES-Blast as templates (Addgene 85146) and by inserting the products into BamHI/NotI-digested pLV-EF1a-IRES-Neo^31^ (Addgene 85139) using Gibson assembly. The stoichiometric ezrin-mRuby3/miRFP680 reporter, in pLV-EF1a-MCS-IRES-Neo (Addgene 85139), consisted of full-length ezrin (PCR-amplified from HT-HUVEC cDNA), C-terminally tagged with mRuby3, followed by a p2A ribosomal skipping sequence^65^ and a cytosolic miRFP680^91^. The sequence encoding miRFP680 was downloaded from Addgene (136557, pmiRFP680-N1), optimized for mammalian expression using a web-based codon-optimization tool (Thermo Fisher Scientific), and synthesized (gBlock, IDT DNA) with a nuclear export signal (LALKLAGLDI)^92^ and p2A sequence appended.

Plasmid constructs generated during this study will be available from Addgene following publication.

### Lentivirus production

Lentivirus was generated in HEK293FT cells as described^31^. Briefly, confluent 10cm dishes of cells were co-transfected with a transfer vector containing the gene of interest (15µg) and the 3^rd^-generation packaging plasmids pMDLg/pRRE (5µg), pRSV-rev (5µg), pCMV-VSVG (5µg) using Lipofectamine2000 (50µl, Thermo Fisher Scientific 11668027), in final 10ml of OptiMEM (Thermo Fisher Scientific 31985070). Viral supernatants were collected at 48h post transfection, 0.22µm-filtered (VWR CA28143-310), concentrated using centrifugal filter units (100 kDa cutoff, MilliporeSigma, UFC910024), aliquoted, and either used immediately or stored at -80°C.

### Cell plating, live-cell, and fixed cell microscopy

Optical 96-well glass-bottom plates (Cellvis P96-1.5H-N) were coated with bovine collagen type I, 31µg/ml in PBS, for 2-4h at 37°C. Cells were plated at densities and for durations prior to fixation or live-cell imaging as specified. For live-cell imaging, cells were overlaid with a CO_2_-independent live-cell imaging solution (LIS) with low autofluorescence, composed of 125mM NaCl, 5mM KCl, 1.5mM MgCl_2_, 1.5mM CaCl_2_, 10mM D-glucose, 20mM HEPES pH 7.4, 1% FBS and 5ng/ml bFGF, and plates were sealed during imaging using aluminum microplate seals (PolarSeal, Thomas Scientific 1152A34). Cells were fixed by adding fixation solution (4% formaldehyde in PBS) at a ratio of 1:1 to culture medium or LIS (final 2% formaldehyde) and incubated for 15min at RT. Following two PBS washes, cells were either stained with WGA-AF647 (2.5µg/ml in PBS, 10min at RT), or permeabilized for immunofluorescence staining using ASBB, a permeabilization/blocking solution (10% FBS, 1% BSA, 0.1% Triton X-100, 0.01% NaN_3_, in PBS) for 30min. Incubation with primary antibodies, diluted in ASBB 1:400, was done either at RT for 2h or at 4°C overnight. Incubation with secondary antibodies, diluted 1:1000 in ASBB, was done at RT for 1h. Fixed cells were imaged overlaid with PBS.

Imaging data were acquired using one of the four livecell imaging systems described below, as indicated in the following table:

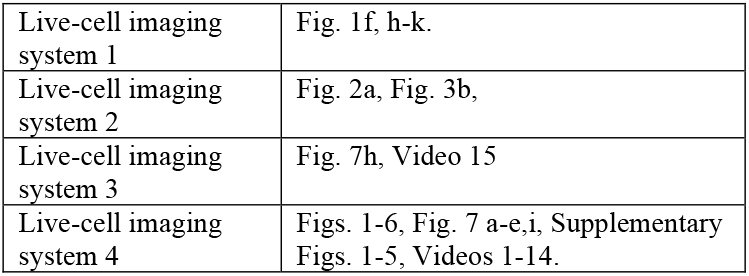

#### Live-cell imaging system 1

Fully automated widefield/Yokogawa spinning disc confocal fluorescence microscope system (Intelligent Imaging Innovations, 3i), built around a Nikon Ti-E stand, equipped with Nikon ‘s Perfect Focus System, a 40x 1.3NA Plan Fluor oil objective, a 3i laser stack (405, 442, 488, 514, 561, 640nm), a broad-range fluorescence light source with integrated excitation filter wheel, (Lambda XL, Sutter), a Yokogawa CSU-W1 scanning head with dual camera port and emission filter wheels, two sCMOS cameras (Andor Zyla 4.2), enclosed by an environmental chamber (Haison), and controlled by SlideBook software (3i).

#### Live-cell imaging system 2

Fully automated fluorescence microscope system (ImageXpress Micro XL, Molecular Devices), equipped with white light LED light source (SOLA, Lumencor), a Zyla 5.5 sCMOS camera (Andor), a 20x 0.75 Plan Apo objective (Nikon), and controlled by MetaXpress software.

#### Live-cell imaging system 3

Fully automated widefield fluorescence microscope system (Nikon), built around a Nikon TI-E stand, equipped with Nikon ‘s Perfect Focus System, a 10x 0.45 NA Plan Apo air, a 20x 0.75 NA Plan Apo air, and a 40x 1.3 NA Plan Fluor oil immersion objective, a liquid light guide-coupled white light LED light source (SOLA SE UV-nIR, Lumencor), excitation and emission filter wheels (Lambda 10-3, Sutter), a sCMOS camera (Prime-BSI, Photometrics), enclosed by a custom-built environmental chamber (Digital Pixel), and controlled using Nikon NIS Elements AR software.

#### Live-cell imaging system 4

Fully automated widefield fluorescence microscope system (Nikon), built around a Nikon TI2-E stand, equipped with Nikon ‘s Perfect Focus System, a 10x 0.45 NA Plan Apo air, a 20x 0.75 NA Plan Apo air, and a 40x 1.3 NA Plan Fluor oil immersion objective (Nikon), a liquid light guide-coupled multispectral LED light source (SpectraX, Lumencor), a dual-camera image splitter (TwinCam, Cairn) with a custom-integrated high-speed emission filter wheel (HS-1025, FLI), two sCMOS cameras (Orca Fusion-BT, Hamamatsu), a triggered device hub (NI-BB, Nikon), enclosed by a custom-built environmental chamber (Digital Pixel), and controlled using Nikon NIS Elements AR software.

### Transient transfection of cDNA, synthetic Rho, Rac, Ccd42 (in)activation using rapamycin-induced heterodimerization

HT-HUVEC expressing RhoB sensor, RaichuEV-Rac, or RaichuEV-Cdc42 were plated in collagen-coated wells of 96-well glass-bottom plates at 15,000 cells/well the day before the transfection. The day of the transfection, the culture medium was replaced with antibiotics-free full growth medium, 80µl per well. Then, 0.2µg of DNA (LynFRB : mCherry-FKBP constructs 5:1 w/w) and 0.25µl of Lipofectamine 2000 (Thermo Fisher Scientific, 11668019), diluted in 20µl OptiMEM (Thermo Fisher Scientific, 31985070), was added as per manufacturer ‘s recommendation. The transfection mix was replaced after 2h with full growth medium. 16-18h later, cells were overlaid with LIS and images captured using live cell imaging system 1, using a 40x 1.3 NA oil objective lens, at 0.33µm/pixel resolution (2×2 binning). FRET, CFP, mCherry channels were acquired sequentially. 60 images were captured at 15s intervals, Rapamycin was added after timepoint 10 at 0.5µM final.

### Image analysis – background subtraction, cell segmentation and cell tracking

Background images for each channel were generated by imaging wells without cells and filled with LIS. Multiple images from multiple wells were averaged and subjected to circular image filtering using MATLAB functions *imfilter* and *fspecial*. To adjust raw images of cells, the median intensity of neighboring pixels from the average background image was subtracted. Cell masks were generated using histogram-based thresholding based on pixel intensity distribution of the YFP-FRET channel. To smooth the cell edge and refine the mask, Gaussian circular averaging filters were applied. To remove small debris trails attached to/touching cells, MATLAB functions *imopen* and *strel* were used to remove edge elements with a radius less than 1 pixel. Subsequent cell trajectories were created using a nearest-neighbor consecutive pairing algorithm as described previously^19^.

### YFP-FRET and CFP channel alignment, FRET/CFP ratio, and fluorophore bleaching correction

Methods for YFP-FRET and CFP channel registration have been described previously^19^. FRET/CFP ratios were computed as per-pixel ratios from background-subtracted, registered, and noise-filtered YFP-FRET and CFP images. By plotting FRET/CFP averaged per field of view over time from untreated samples, a bleaching correction curve was generated, displaying exponential-like decay. Assuming FRET/CFP across multiple untreated cells remained constant throughout an experiment, the FRET ratio array for each time frame was divided against the curve, normalizing it.

### Image analysis – mapping cell-edge velocity and RhoGTPase activities

Cell edge velocities and RhoGTPase activities in peripheral coordinate windows were analyzed as described previously^19^. Briefly, the cell edge of segmented cells was divided into 900 equally spaced coordinate windows. For each coordinate window, velocity vectors were formed by the dot product between a unit vector normal to the cell edge and the displacement vector from frame to frame. Coordinate windows were then binned into groups of 5, forming 180 averaged coordinate windows, each with their own velocity vector. Coordinate windows had an adjustable depth parameter, including 0.98, 2.0, 3.3, 4.9, 6.5, and 8.1µm. FRET/CFP or any expressed protein ‘s value for each coordinate window was the average value of all pixels assigned to the window. To remove stochastic noise from edge velocity and FRET activity data, the MATLAB function *ndnanfilter* was applied.

### Image analysis – thresholding protrusions and retractions

Cell edge velocity maps were first thresholded either above 3.9µm/min, creating a map of only protrusive activity, or below -3.9µm/min to depict retractive activity. Using MATLAB ‘s *bwareaopen* function, we then applied a minimum size threshold of 25 pixels to each map, leaving distinct protrusion and retraction events above the size threshold. For each event, data such as area size, pixel IDs, time range, cell edge coordinate range, average edge velocity, average FRET activity were recorded.

### Cross-correlation analysis

All cross-correlation analyses were performed using MATLAB’s *xcorr* function. Cell edge velocity and protein activation arrays were shifted to a Standard Normal ∼ (0,1) distribution, using the Central Limit Theorem. To avoid a built-in time lag when comparing FRET/CFP to edge velocity at a given frame x, edge velocities from frame x-1 ⟶ x and x ⟶ x+1 were averaged. Then, cross-correlation was computed for each window coordinate vector of all time points, yielding 180 correlation vectors. These were averaged to form one cross-correlation vector per cell. The overall cross correlation for a given experimental condition was obtained by averaging the vectors from all cells analyzed.

### Edge velocity vs. RhoGTPase activity buildup plots

Previously generated edge velocity accompanying FRET/CFP arrays from sample cells were loaded and processed using MATLAB. To avoid bias, only edge velocity maps were visualized when choosing protrusion-retraction transition events. Rectangular regions of interest undergoing protrusion retraction cycling were identified. A region of interest spanning 15 coordinate windows or 8.3% of the cell perimeter was created. Only events with > 20 frames or 8.3min of averaged positive edge velocity preceding the transition event were used. The exact transition point from protrusion to retraction was identified as the first frame x with (i) negative acceleration, (ii) positive edge velocity at frame x-1, and negative edge velocity at frame x. Between frame x-1 and x, the frame with lowest absolute-valued edge velocity was used as the transition point. The length of the rectangular ROI was 41 frames, using ± 20 frames before and after the transition frame (Supplementary Fig. 2 a,b). Edge velocity and FRET/CFP at a chosen edge depth were plotted on the same graph (Supplementary Fig. 2c).

### Image analysis – ezrin-mRuby3/miRFP680 ratio calculation

Ezrin-mRuby3 and miRFP680 image channels were background corrected and segmented using the YFP-FRET derived cell mask. Circular averaging image filters were applied to each channel (*ndanfilter* and *fspecial* in MATLAB). Each channel was normalized to its bleaching curve. Then, the per-pixel ezrin-mRuby3/miRFP680 ratio was computed. The 1^st^ and 99^th^ percentile values of the ratio were calculated, and used as the lower and upper limits, i.e., any ratio value less than the 1st percentile was rounded up to the first percentile, and any values above the 99^th^ percentile was rounded down to the 99^th^ percentile. This was to avoid outlier extreme ratio values due to stochastic noise.

### Quantitative immunofluorescence

Fixed cells in 96-well glass bottom dishes were stained with 1:400 diluted anti-pERM antibody, (1:1000 diluted AF568-conugated secondary antibody), 1:200 diluted AF488-conjugated phalloidin, and 1:10,000 diluted Hoechst. 16 images were captured per well, using a 20x 0.75 NA air objective at 0.33µm/pixel resolution. Images with visible staining or imaging artifacts were discarded, the remaining analysis was fully automated. Cell nuclei were detected based on the Hoechst images using a previously described MATLAB routine^93^. The nuclei in the resulting mask were dilated by 7 pixels (2.3µm) and the resulting image regions used to determine per-cell pERM signals, using background-subtracted pERM images.

### RhoGTPase activity response to rapamycin-induced Rho, Rac and Cdc42 (in)activation

Images were processed and raw FRET/CFP ratios were computed as described above. An interactive custom MATLAB routine was then used to manually identify cells co-expressing mCherry-FKBP-tagged RhoGTPase regulatory domains and Lyn-FRB, based on the presence of mCherry fluorescence. ROIs were drawn within cells identified this way and the corresponding raw FRET/CFP time-courses automatically computed. For controls, ROIs were drawn in cells without mCherry expression. All timecourses were first individually normalized to the average FRET/CFP of the timepoints prior to rapamycin addition. Then, the time-courses of mCherry-expressing cells were normalized by averaged control time-courses, before averaging per condition, across two biological replicates.

### RhoGTPase activity responses to drug additions

Cells were plated 3-4h before imaging in collagen-coated glassbottom 96 well plates at a density of 2,000 cells/well. Full growth medium was replaced with LIS 1h prior to imaging. Time lapse sequences were acquired at 20x magnification, 2×2 binning, 0.65µm/pixel, at 1min intervals for 60min. Drugs diluted in LIS and LIS only (control) were added to wells after indicated times. Image data was processed as described above and a pixel-wise YFP-FRET/CFP calculation was performed. For each field of view, average Rho-FRET activity per frame was calculated and normalized (1) to its own FRET average in the frames pre-drug addition, and (2) to the mean control FRET response from all control FOVs. These normalizations scaled each FRET response to 1 during the pre-drug addition period and accounted for any increase in FRET activity due to the addition of control LIS.

### Cell shape analysis

HT-HUVEC were seeded at 750 cells per well on collagen-coated 96-well glass-bottom plates in either full growth medium (control) or full growth medium supplemented with either Cpd31 (5µM), Y27632 (10µM), or both Cpd31 (5µM) and Y27632 (10µM). Cells were incubated for 2h at 37°C, before fixation with 2% formaldehyde in PBS and staining with AF647-conjugated wheat germ agglutinin (WGA, 2.5µg/ml in PBS, 10min at RT) as a membrane marker and Hoechst (1:10,000) to stain the nuclei. Images were then acquired using live-cell imaging system 4, acquiring 16 sites per well across 30-40 wells at 20x magnification with a pixel size of 0.33µm. Image processing was performed using custom MATLAB code. Objects (cells) were automatically detected and cropped from each image. To exclude cell debris and other particles, only objects with an area greater than 3,000 pixels were retained. Object properties (centroid coordinates, area, eccentricity, solidity, major axis length, minor axis length, perimeter, major axis orientation and mean intensity) were then extracted from each cropped image, as defined by MATLAB ‘s built-in *regionprops* function. Solidity is defined as the ratio of the area occupied by the cell over the area of the convex hull of the cell shape. Eccentricity is defined as the ratio between the inter-focal distance and the major axis of an ellipsoid. The number of objects in the nuclear and cell masks were used to discard objects without a nucleus (i.e., large cell debris), and cases with more than one nucleus but only one object (i.e., cells touching each other and binucleate cells).

### Cell area and blebbing quantification

RhoB sensor-expressing HT-HUVEC were seeded at 1,000 cells per well in collagen-coated 96-well glass-bottom plates and incubated in full growth medium for 3-4h before replacing the full growth medium by LIS. At specified times during image acquisition, LIS (control), nocodazole (15µM), both nocodazole (15µM) and Cpd31 (5µM) or Cpd31 (5µM) were administered to the corresponding wells. Image series were acquired using live-cell imaging system 3 or 4 at 1min intervals over 135min in the YFP channel using 10x magnification and 0.65µm/pixel resolution. Image analysis was performed using custom MATLAB code. The image series were registered using the hardware stage position data and the background illumination profile was corrected using images from wells left without cells. Cell segmentation was then performed using a local minima histogram-based segmentation algorithm to extract cell occupancy area per field of view per frame. Then, cells were manually counted and classified as either blebbing or spread from images taken from one frame before drug addition and at the end of the time series.

### Statistical testing and reproducibility

For comparison of two groups, the non-parametric Whitney-Mann U rank sum test was used. For comparison of multiple groups, one-way ANOVA followed by the Tukey-Cramer ‘s pairwise comparisons were performed. All data shown are from multiple biological and technical replicates, as specified in the figure legends. Statistical analyses were performed using MATLAB or Graphpad Prism.

### Code and Data Availability

Data and code are available from the authors upon reasonable request.

## Video legends

**Video 1**

HT-HUVEC stably expressing the RhoB sensor was imaged at 25sec intervals. Video corresponds to Fig. 1a. YFP-FRET channel is shown, displayed at 15fps. Scale bar, 20µm.

**Video 2**

Random motility of a HT-HUVEC stably expressing RaichuEV-Rac. *(Left)* Probe localization, *(right)*, FRET/CFP, scaling [0.7 1.3]. Acquired at 25sec intervals, displayed at 15fps. Scale bar, 20μm.

**Video 3**

Random motility of a HT-HUVEC stably expressing RaichuEV-Cdc42. *(Left)* Probe localization, *(right)*, FRET/CFP, scaling [0.7 1.3]. Acquired at 25sec intervals, displayed at 15fps. Scale bar, 20μm.

**Video 4**

Random motility of a HT-HUVEC stably expressing RhoB sensor. *(Left)* Probe localization, *(right)*, FRET/CFP, scaling [0.7 of 25 pixels to each map, leaving distinct protrusion and retraction events above the size threshold. For each event, data such as area size, pixel IDs, time range, cell edge coordinate range, average edge velocity, average FRET activity were recorded.

**Video 5**

Random motility of a HT-HUVEC stably expressing DORA-RhoA. *(Left)* Probe localization, *(right)*, FRET/CFP, scaling [0.7 1.3]. Acquired at 25sec intervals, displayed at 15fps. Scale bar, 20μm.

**Video 6**

Random motility of a HT-HUVEC stably expressing RhoA2G. *(Left)* Probe localization, *(right)*, FRET/CFP, scaling [0.7 1.3]. Acquired at 25sec intervals, displayed at 15fps. Scale bar, 20μm.

**Video 7**

HT-HUVEC stably expressing RhoB sensor during initial adhesion formation, spreading, and random motility. Retraction example in Fig. 3a,b occurs between t = 20 and 30min. *(Left)* Probe localization, *(right)*, FRET/CFP, scaling [0.7 1.3]. Acquired at 25sec intervals, displayed at 15fps. Scale bar, 25µm.

**Video 8**

HT-HUVEC stably expressing RhoB sensor exhibiting oscillatory RhoGTPase activity, characteristic of repeated protrusion-retraction cycles. *(Left)* Probe localization, *(right)*, FRET/CFP, scaling [0.7 1.3]. Acquired at 25sec intervals, displayed at 30fps. Scale bar, 20µm.

**Video 9**

HT-HUVEC stably expressing RhoB sensor exhibiting a propagating wave of paired elevated RhoB activity and edge retraction. *(Left)* Probe localization, *(right)*, FRET/CFP, scaling [0.7 1.3]. Acquired at 25sec intervals, displayed at 30fps. Scale bar, 20µm.

**Video 10**

HT-HUVEC stably co-expressing RhoB sensor and mRuby3-MLC undergoing random motility. Retraction example in Fig. 4a occurs between t=0 min and t=23min. *(Left)* RhoB sensor FRET/CFP, scaling [0.7 1.3], *(right)* mRuby3-MLC. Acquired at 15sec intervals, displayed at 30fps. Scale bar, 20µm.

**Video 11**

HT-HUVEC stably co-expressing RhoB sensor and mRuby3-MLC, treated with 20µM ML7 at 5min. Channels from left to right: RhoB sensor localization, Rho sensor FRET/CFP [0.5 1.5], and mRuby3-MLC. Acquired at 1 min intervals, displayed at 15fps. Scale bar, 20µm.

**Video 12**.

HT-HUVEC stably co-expressing RhoB sensor and mRuby3-MLC, treated with 20µM Y27632 at 5 min. Channels from left to right: RhoB sensor localization, Rho sensor FRET/CFP [0.5 1.5], and mRuby3-MLC. Acquired at 1min intervals, displayed at 15fps. Scale bar, 20µm.

**Video 13**

HT-HUVEC stably co-expressing RhoB sensor and mRuby3-MLC. *(Left)* RhoB sensor FRET/CFP, scaling [0.7 1.3], *(center)* mRuby3-MLC. Video displays movement of cell shown in Fig. 5a during 14min preceding fixation and anti-pERM immunostaining. (*right*) The same cell after anti-pERM immunostaining. Elevated pERM signal colocalizes with elevated RhoB sensor activity in retraction. Acquired at 1min intervals, displayed at 5fps. Scale bar, 20µm.

**Video 14**

HT-HUVEC stably coexpressing RhoB sensor and ezrin-mRuby3-P2A-miRFP680-NES during random motility. Channels from left to right: RhoB sensor localization, RhoB sensor FRET/CFP, scaling [0.7 1.3], ezrin-mRuby3/miRFP680 ratio, scaling [0.5 1.5]. Acquired at 25sec intervals, displayed at 15fps. Scale bar 20µm.

**Video 15**

HT-HUVEC stably expressing RhoB sensor were treated as depicted in Fig. 7h, video corresponds to Fig. 7g. *(Left)* control, *(center left)* Nocodazole, 15µM, added at 15min and 45min. *(center right)* Nocodazole, 15µM, and Cpd31, 5µM, added at 15min. *(Right)* Nocodazole, 15µM, added at 15min, Cpd 31, 5µM, added at 45min. RhoB sensor localization is shown. Acquired at 1min intervals, displayed at 15fps. Scale bar, 100µm.

## Acknowledgements

We thank Sean Collins for sharing analysis code, Olivier Pertz and Yi Wu for providing constructs. We are grateful to Aparna Suvrathan, Greg Emery, and Tobias Meyer for helpful discussions and comments on the manuscript. This work was supported by an NSERC Discovery Grant (RGPIN-2018-05831), a CIHR Project Grant (PJT-165932), a CFI-JELF Equipment Grant (37549), an FRQNT Lab Startup Grant (FRQ-NT NC-298218), and McGill Startup Funding to A.H.. S.M.B. was supported by an NSERC Undergraduate Summer Research Award and an NSERC Canada Graduate Scholarship Masters Award, R.A.M.R by an FRQNT Doctoral Training Scholarship, and R.S. by an. NSERC Undergraduate Summer Research Award. Some of the image data collection was performed in the McGill University Advanced BioImaging Facility (ABIF), RRID; SCR_017697

## Author contributions

S.M. and A.H. conceived the project. Data collection and analysis: Figs. 1-3 S.M. and A.H., Fig. 4-5 S.M., Fig. 6 A.H., Fig. 7 R.A.M.R., A.H., S.M., R.S. and M.U. Supplementary Fig. 1-4 S.M., Supplementary Fig. 5 N.E.B. and R.A.M.R. The ezrin reporter used in Fig. 5 was developed by Q.L. Analysis code development: A.H., S.M., and R.A.M.R. S.M., A.H. and RMR wrote the paper, with input from all authors.

