## Supplementary Figures 1-5 for "Excitable Rho dynamics drive cell contractions by sequentially inducing ERM protein-mediated actin-membrane attachment and actomyosin contractility"

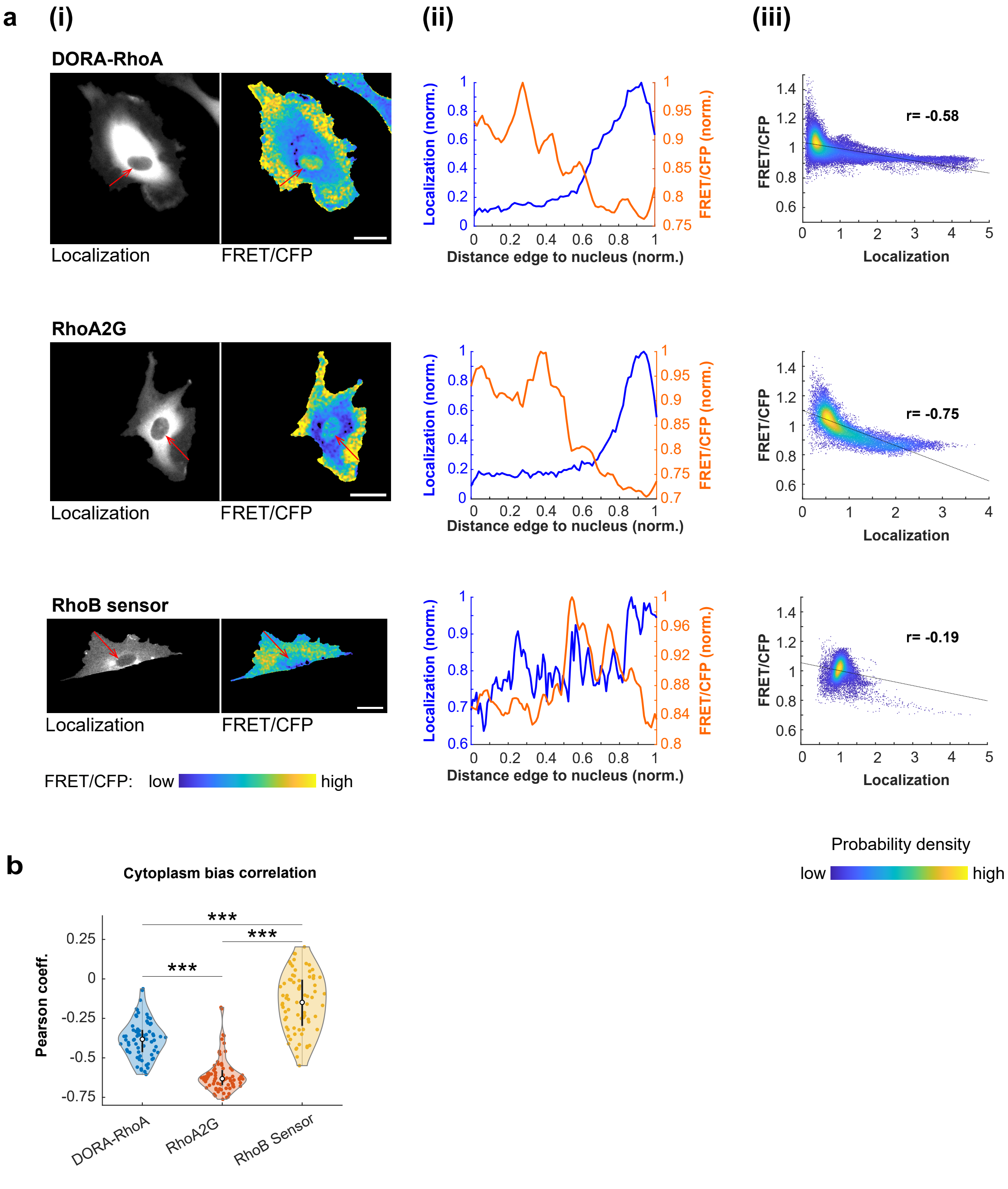


**Supplementary Figure 1. Localization/cell geometry-imposed bias on activity reported by the Rho FRET probes DORA-RhoA, RhoA2G, and RhoB sensor**

**(a) (i)** Side-by-side comparisons of FRET probe localizations *(left)* and FRET/CFP ratios *(right)* for DORA-RhoA, RhoA2G, and RhoB sensor. Scale bars, 25µm. Quantifications along the red arrows, manually drawn from the cell edge to the nucleus, are shown in (ii). **(ii)** Line plots from the images in (i) depicting normalized localization compared to normalized FRET/CFP ratio. Both were normalized to the maximum value along the line profiles. **(iii)** Per-pixel correlation between localization intensity and FRET/CFP ratio of the two images in (i), with FRET/CFP plotted as a function of normalized localization signal. Pearson R correlation coefficients were calculated using MATLAB *corrcoeff* function, and line of best fit was plotted using MATLAB *polyfit* function. **(b)** Compiled Pearson R correlation coefficients for DORA-RhoA, RhoA2G, and RhoB sensor. Individual frames from time-lapse sequences from biological replicates were chosen at random and the correlation coefficient between normalized localization and FRET/CFP were calculated. DORA-RhoA: n= 69 frames from 23 cells, 2 independent trials. RhoA2G: n=75 frames from 25 cells, 2 biological replicates. DORA-RhoB: n=75 frames from 25 cells, 3 biological replicates. The bolded circle in violin plot shows dataset median, and bolded black lines show 25th and 75th percentiles. ****p<0.001*, one-way ANOVA/Tukey-Cramer.

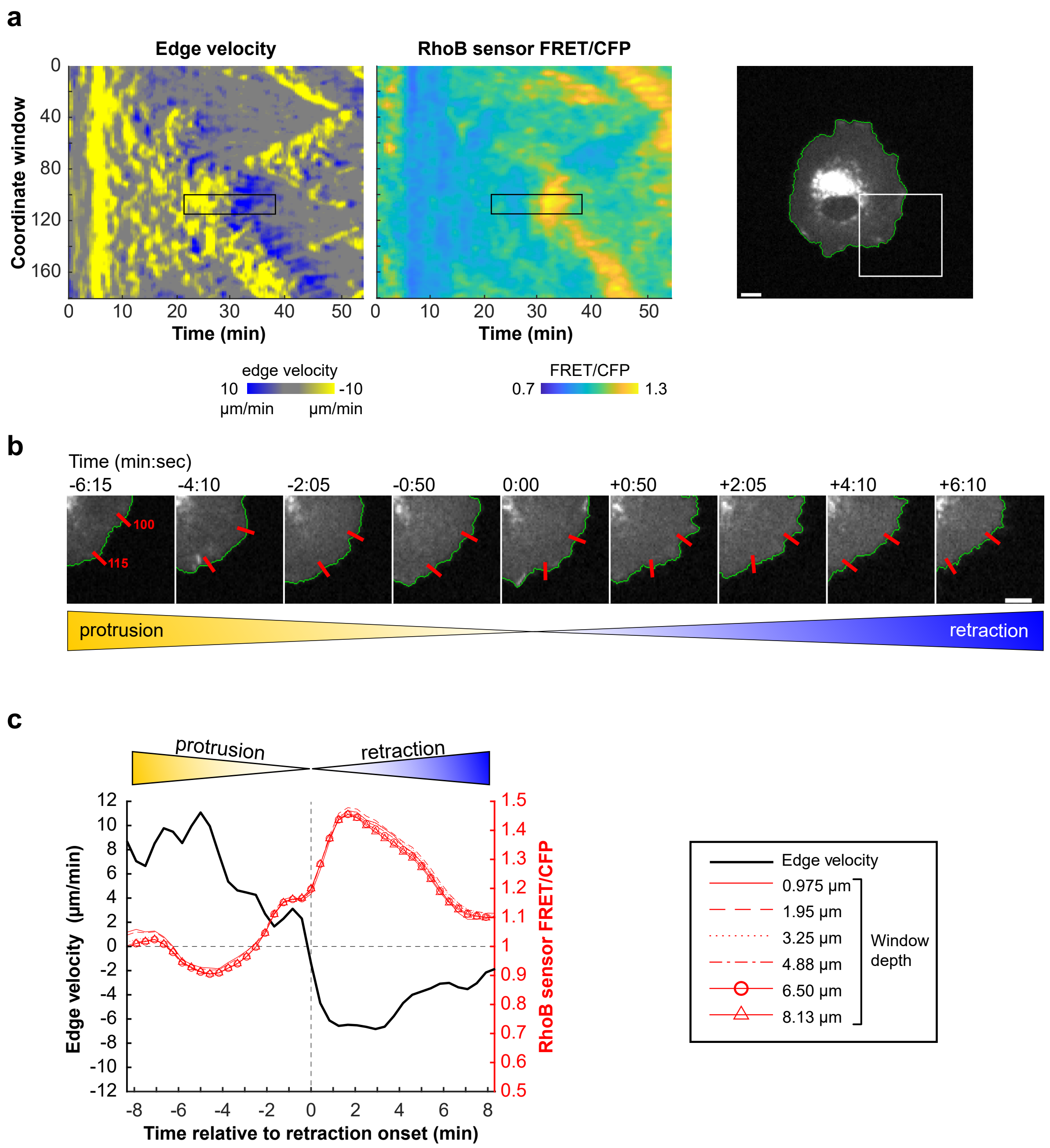


**Supplementary Figure 2. Edge velocity-Rho activity buildup analysis**

**(a)** Edge velocity and Rho sensor FRET/CFP spatiotemporal heatmaps with region of interest outlined by a black rectangle. Black rectangle is 15 coordinate windows wide, approx. 8.33% of the cell perimeter, spanning a duration of 20min. General region of analysis shown by white rectangle overlaid on cell outline on the right. Scale bar, 10 μm. **(b)** Time-lapse illustrating protrusion-retraction transition. Region of interest between coordinate windows 100 and 115 depicted by red ticks. Scale bar, 10μm. **(c)** Plot for the region of interest from (a) and (b) comparing average edge velocity to average RhoB sensor FRET/CFP activity per timepoint. Time=0 is placed at retraction onset, i.e., the transition from positive to negative edge velocity. RhoB sensor FRET/CFP per timepoint for varying window depths are plotted with different line styles.

**a**

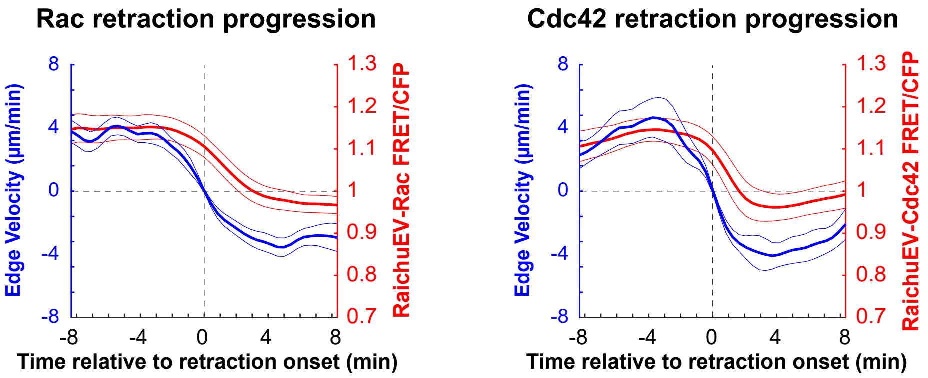


**Supplementary Figure 3. Rac and Cdc42 activity during protrusion-retraction transitions**
**(a)** Edge velocity and RaichuEV-Rac (left) or RaichuEV-Cdc42 (right) FRET/CFP buildup plots displaying FRET/CFP at edge depths of 1.95µm during protrusion-retraction transitions. Time = 0 denotes retraction onset. Dataset means are bolded, bordered by ± 95% CI of the mean. RaichuEV-Rac: n =29 events, 4 biological replicates, RaichuEV-Cdc42: n =17 events, 3 biological replicates.

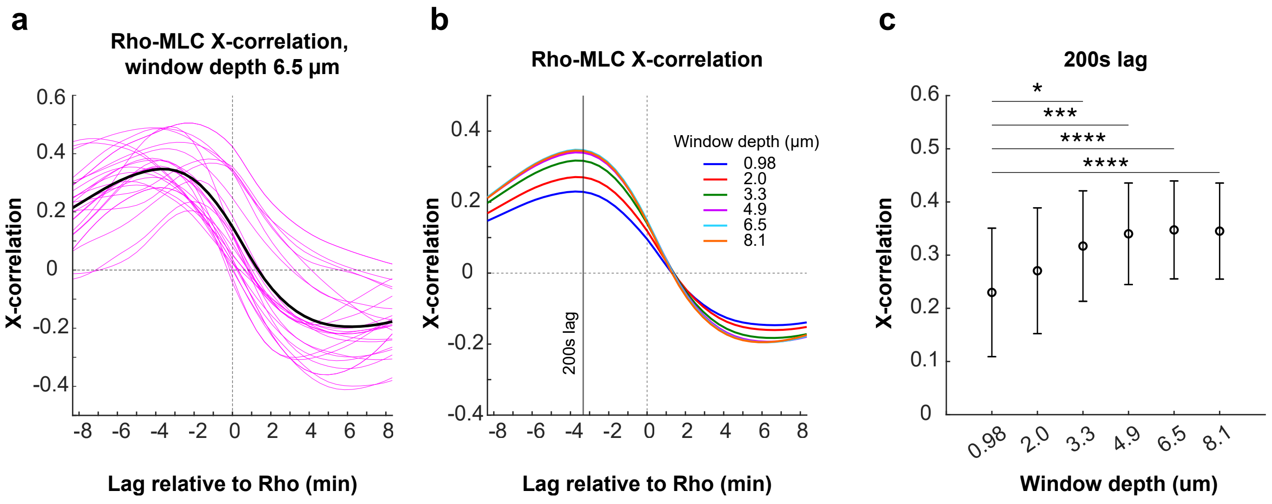


**Supplementary Figure 4. Cross-correlation between RhoB sensor activity and mRuby3-MLC signal**

**(a)** Cross correlation between RhoB sensor FRET/CFP and mRuby3-MLnal at an edge depth of 6.5µm. Pink lines represent results from individual cells, with the mean correlation bolded in black. n =27 cells from two biological replicates. **(b)** Average cross correlation between RhoB sensor FRET/CFP and mRuby3-MLC at window depths varying form 0.98 to 8.1µm. 200 s lag, where all traces reach their maximum, is indicated by a vertical line. n=27 cells from two biological replicates. **(c)** Comparison of average correlation-coefficient at a lag of 200s for all edge depths tested, i.e., a quantification of the difference of the traces in (b) at 200s lag. n=27 cells from two biological replicates. **p<0.05, ***p<0.001, ****p<0.0001*, one-way ANOVA/Tukey-Cramer.

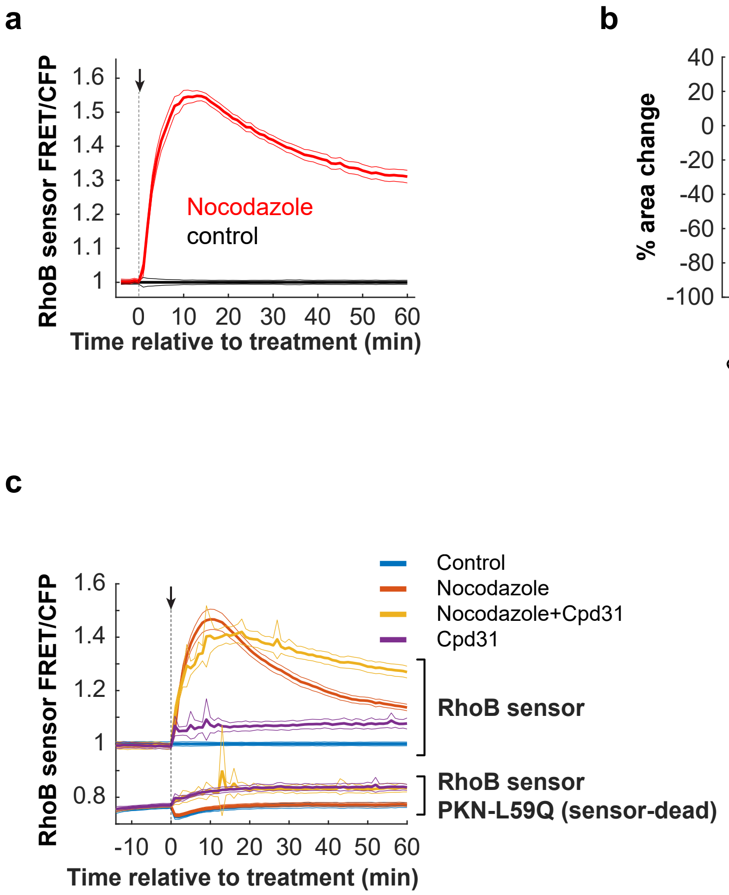


**Supplementary Figure 5. Additional RhoB sensor validations**

**(a)** RhoB sensor activity response to acute nocodazole addition (10μM). Treated in red, normalized to control addition in black. Bolded lines denote mean, bordered by ± 95% CI. n=36 treated and control fields of view (FOV) each were analyzed, from 4 biological replicates. **(b)** Percent area changes of cells in control vs. nocodazole treated FOV, measured 30min post nocodazole addition. n=36 treated and control FOVs, from 4 biological replicates. ****p<0.001*, Mann-Whitney U-test. **(c)** Responses of RhoB sensor and of a sensor-dead version of the RhoB sensor with RhoGTP binding-deficient PKN (PKN-L59Q) to control treatment or to treatment with nocodazole (10μM), Cpd31 (5μM), or nocodazole & Cpd31. All data were normalized to control-treated RhoB sensor responses. The presence of Cpd31 did not prevent nocodazole-induced Rho activation. The sensor-dead version of RhoB sensor PKN-L59Q showed minimal responses to all treatments. For each time-lapse sequence, the per frame average ratiometric FRET activity was calculated and the means of 16 time-lapse sequences (FOV) per condition are shown (bold), along with the 95% confidence intervals (thin). Four biological replicates, n=16 FOV for each condition.
